# LY6E impairs coronavirus fusion and confers immune control of viral disease

**DOI:** 10.1101/2020.03.05.979260

**Authors:** Stephanie Pfaender, Katrina B. Mar, Eleftherios Michailidis, Annika Kratzel, Dagny Hirt, Philip V’kovski, Wenchun Fan, Nadine Ebert, Hanspeter Stalder, Hannah Kleine-Weber, Markus Hoffmann, H. Heinrich Hoffmann, Mohsan Saeed, Ronald Dijkman, Eike Steinmann, Mary Wight-Carter, Natasha W. Hanners, Stefan Pöhlmann, Tom Gallagher, Daniel Todt, Gert Zimmer, Charles M. Rice, John W. Schoggins, Volker Thiel

**Author notes:** equally contributing authors (Stephanie Pfaender, Katrina B. Mar). equal senior authorship.

## Abstract

Zoonotic coronaviruses (CoVs) are significant threats to global health, as exemplified by the recent emergence of severe acute respiratory syndrome coronavirus 2 (SARS-CoV-2)^1^. Host immune responses to CoV are complex and regulated in part through antiviral interferons. However, the interferon-stimulated gene products that inhibit CoV are not well characterized^2^. Here, we show that interferon-inducible lymphocyte antigen 6 complex, locus E (LY6E) potently restricts cellular infection by multiple CoVs, including SARS-CoV, SARS-CoV-2, and Middle East respiratory syndrome coronavirus (MERS-CoV). Mechanistic studies revealed that LY6E inhibits CoV entry into cells by interfering with spike protein-mediated membrane fusion. Importantly, mice lacking Ly6e in hematopoietic cells were highly susceptible to murine CoV infection. Exacerbated viral pathogenesis in Ly6e knockout mice was accompanied by loss of hepatic and splenic immune cells and reduction in global antiviral gene pathways. Accordingly, we found that Ly6e directly protects primary B cells and dendritic cells from murine CoV infection. Our results demonstrate that LY6E is a critical antiviral immune effector that controls CoV infection and pathogenesis. These findings advance our understanding of immune-mediated control of CoV *in vitro* and *in vivo*, knowledge that could help inform strategies to combat infection by emerging CoV.

---

Coronaviruses (CoVs) are enveloped RNA viruses with broad host tropism and documented capability to cross the species barrier to infect humans^3^. The mammalian innate immune response controls CoV infection partly through the action of interferons (IFNs)^2,4,5^. IFNs inhibit viral infection by inducing hundreds of genes, many of which encode antiviral effectors^6^. The interferon-stimulated genes (ISGs) that inhibit CoV infection are not well defined^2^. To address this knowledge gap, we screened our published library of >350 human ISG cDNAs^7,8^ in human hepatoma cells for antiviral activity against the endemic human CoV 229E (HCoV-229E) (**Extended Data Table 1, Extended Data Table 2**). The ISG lymphocyte antigen 6 complex, locus E (LY6E) strikingly inhibited HCoV-229E infection (**Fig. 1a-b**, **Extended Data Fig. 1a**). This was unexpected since we and others previously reported that LY6E enhances cellular infection by influenza A virus, flaviviruses, and HIV-1^9–11^, with HIV-1 being an exception as opposing roles of LY6E have been described^12^. To verify this result, we generated stable cell lines ectopically expressing LY6E and infected cells with diverse CoVs. In addition to HCoV-229E (**Fig. 1c**), LY6E significantly inhibited infection by HCoV-OC43, MERS-CoV, SARS-CoV, and the recently emerged SARS-CoV-2 (**Fig. 1d-g**). The antiviral effect of LY6E extended beyond human CoVs and blocked infection by the murine CoV murine hepatitis virus (MHV) (**Fig. 1h**). Given the zoonotic origin of SARS-CoV and MERS-CoV, both suspected to originate from bats, we next tested the antiviral potential of select LY6E orthologues. An alignment of LY6E orthologues of human, rhesus macaque, mouse, bat, and camel origin revealed strong conservation across species (**Extended Data Fig. 1b-c**). All LY6E orthologues inhibited HCoV-229E (**Extended Data Fig. 1d**). Recent surveillance of Kenyan camels, the main viral reservoir for MERS-CoV, indicates active circulation of the zoonotic CoV^13^. Human and camel LY6E restricted MERS-CoV infection (**Extended Data Fig. 1e**). To test the role of endogenous LY6E in controlling CoV infections, we used CRISPR-Cas9 gene editing to ablate LY6E expression in human lung adenocarcinoma cells (**Fig. 1i**). This cell line was selected as human hepatoma cells do not express LY6E irrespective of IFN treatment^9^. LY6E knockout (KO) cells were more susceptible to HCoV-229E (**Fig. 1j**). Reconstitution with a CRISPR-resistant LY6E variant (CR LY6E) restored antiviral activity in KO cells (**Fig. 1k-l**). Strikingly, the LY6E-mediated restriction in hepatoma cells was largely specific to CoV. Among 10 genetically divergent viruses tested, only hepatitis C virus was also restricted (**Extended Data Fig. 1f**).

**Figure 1.**
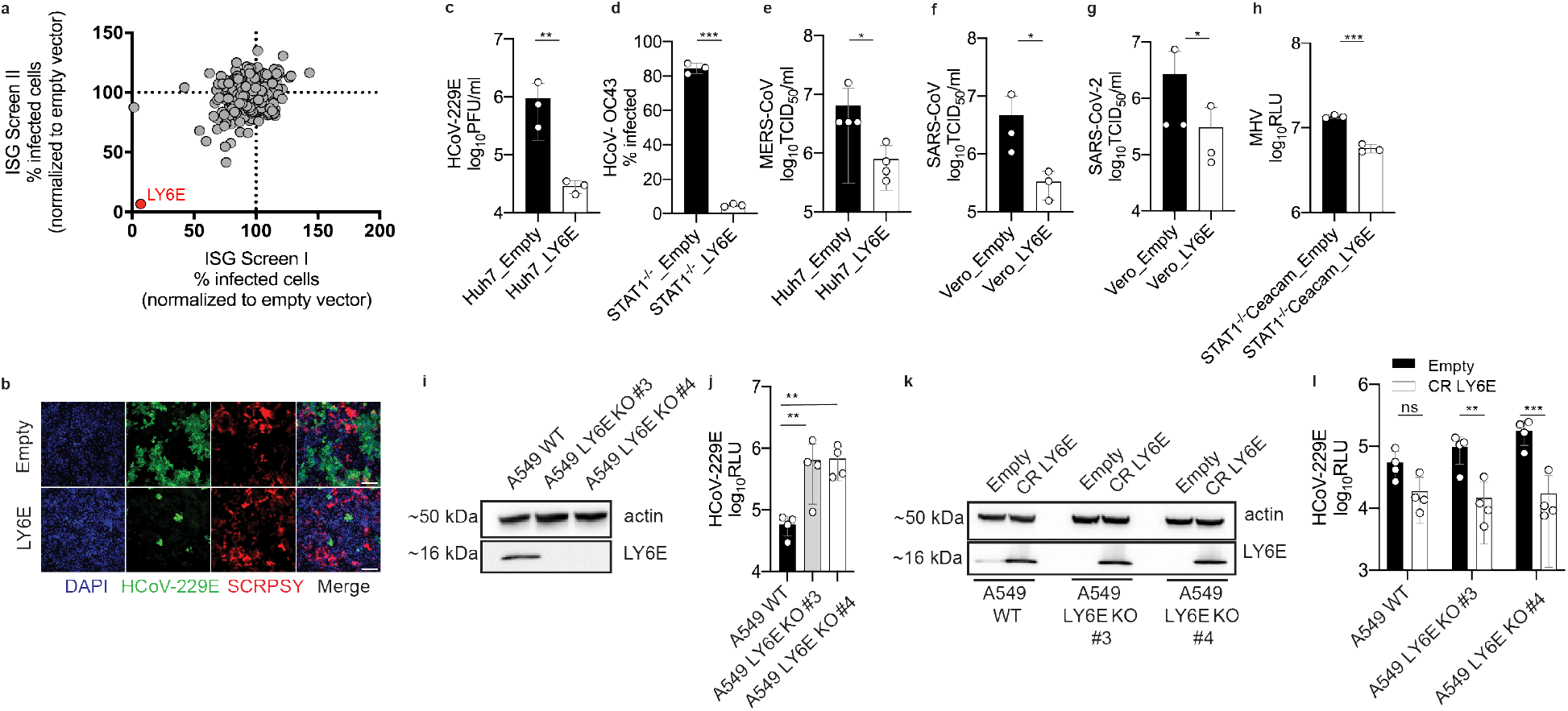
ISG screen identifies LY6E as a potent coronavirus restriction factor. **a**, Duplicate screens depicting HCoV-229E infection (24 hours post-infection) in Huh7 cells ectopically expressing ISGs. **b**, HCoV-229E infection of Huh7 cells expressing LY6E or control vector (Empty). Blue: DAPI, green: HCoV-229E N protein, red: TagRFP encoded in SCRPSY vector. **c-h**, Effect of LY6E expression on diverse coronaviruses. Stably transduced cells were infected with HCoV-229E (**c**), HCoV-OC43 (**d**), MERS-CoV (**e**), SARS-CoV (**f**), SARS-CoV-2 (**g**), MHV-Gluc (**h**). **i**, Western blot of WT A549 or two LY6E KO clones (#3, #4). **j**, HCoV-229E infection of WT or LY6E KO A549. **k**, Western blot of WT or LY6E KO A549 reconstituted with CRISPR-resistant LY6E (CR LY6E) or control vector (Empty). **l**, HCoV-229E infection of CR LY6E-reconstituted WT or LY6E KO A549. Viral replication readouts included: titer determination by plaque assay (**c**), infectivity by nucleoprotein staining and quantitation by flow cytometry (**d**), limiting dilution assay (**e,f,g**), luciferase activity shown as relative light units (RLU) (**h,j,l**). Data represent average of independent biological replicates, n=3 (**c-d,f-h**), n=4 (**e**,**j,l**). In **b**, scale bars are 200 µM. Statistical significance was determined by unpaired student’s t-test with Welch’s correction (**c-d**, **f-g**), one-tailed Mann-Whitney U test (**e**), ratio paired student’s t-test (**g**), 2way ANOVA followed by Sidak’s or Dunnett’s multiple comparison test (**j**,**l**). Error bars: SD. *P* values: **c**, ** p=0.0088; **d**, *** p=0.0001; **e**, * p=0.0143; **f**, * p=0.0286; **g**, * p=0.0101; **h**, *** p=0.0010; **j**, ** p=0.0073, p=0.0033; **l**, ns=0.0740, ** p=0.0013, *** p=0.0001.

LY6E is a member of the LY6/uPAR family of GPI-anchored proteins, and is implicated in diverse cellular processes^14^. To evaluate the specificity of LY6E for inhibiting CoV, we ectopically expressed select LY6/uPAR proteins^9^ and infected with HCoV-229E. Only LY6E inhibited CoV infection (**Extended Data Fig. 1g**). We previously showed that a conserved residue (L36) is required for LY6E-mediated enhancement of influenza A virus^9^. L36 was also required to restrict HCoV-229E infection (**Extended Data Fig. 1h**). These data suggest a conserved functional domain that is responsible for both the virus-enhancing and -restricting phenotypes.

Next, we investigated the effect of LY6E on specific steps of the virus replication cycle. First, we tested whether LY6E restricts HCoV-229E attachment to the cell surface and observed no effect (**Extended Data Fig. 2a**). In accordance, LY6E had no effect on surface expression of CD13, the specific receptor for HCoV-229E (**Extended Data Fig. 2b**). To test whether LY6E restricts viral entry, we used a vesicular stomatitis virus (VSV) pseudoparticle (pp) system bearing CoV spike (S) proteins. Strikingly, LY6E significantly inhibited VSVpp entry mediated by S proteins from HCoV-229E (**Fig. 2a**), MERS-CoV (**Fig. 2b**), SARS-CoV (**Fig. 2c**), and SARS-CoV-2 (**Fig. 2d**). Entry mediated by VSV glycoprotein G was only marginally impacted by LY6E (**Extended Data Fig. 2c, d**). To test whether LY6E interferes with the fusion of viral and cellular membranes, we performed a syncytia formation assay using propagation competent VSV pseudoviruses co-expressing CoV S protein and a GFP reporter (VSV*ΔG(CoV S)) (**Extended Data Fig. 3a**). LY6E potently inhibited CoV S protein-mediated syncytia formation (**Fig. 2e-f**). To determine whether LY6E impairs S protein expression or maturation, we also performed a heterologous syncytia formation assay. CoV-resistant hamster cells infected with trans-complemented GFP reporter VSV*ΔG(CoV) were co-cultured with susceptible target cells that ectopically expressed LY6E. LY6E again significantly blocked syncytia formation, demonstrating that LY6E inhibits S protein-mediated fusion and not S protein expression or maturation (**Extended Data Fig. 3b-c**). As membrane fusion can occur without syncytia formation, we also performed a quantitative fusion assay in which mixing of cell contents results in complementation of split luciferase (**Fig. 2g**). Cells co-transfected with plasmids encoding CoV S proteins and split luciferase 1-7 were mixed with susceptible target cells co-expressing LY6E or vector control and split luciferase 8-11. LY6E reduced CoV S-mediated cell-cell fusion, indicating that LY6E blocks fusion of viral and cellular membranes (**Fig. 2h**). Time-course experiments revealed that LY6E had no effect on HCoV-229E translation, replication, assembly, or release (**Extended Data Fig. 2e-i**). Collectively, our results demonstrate that LY6E specifically inhibits CoV S protein-mediated membrane fusion, which is required for viral entry. We next addressed whether LY6E modulates proteolytic activation of the S protein. Upon S protein-mediated binding to the respective receptor, host proteases cleave the S protein and make it fusion competent for “early entry” at the cell surface or “late entry” in endosomes^15,16^. Pharmacological inhibition of cell surface and endosomal proteases had no effect on LY6E-mediated restriction of CoV infection (**Extended Data Fig. 4a-c**). Next, we addressed whether LY6E directly interferes with the proteolytic activation of S protein. Mutation of S1/S2 and S2′ in the S protein prevents activation (**Extended Data Fig. 4d**)^16,17^. Ectopic LY6E expression had no effect on MERS-CoV S cleavage or infection by cleavage-resistant MERS-CoV pp (**Extended Data Fig. 4e-g**). Collectively, our data indicate that restriction of viral fusion is independent of S protein activation.

**Figure 2.**
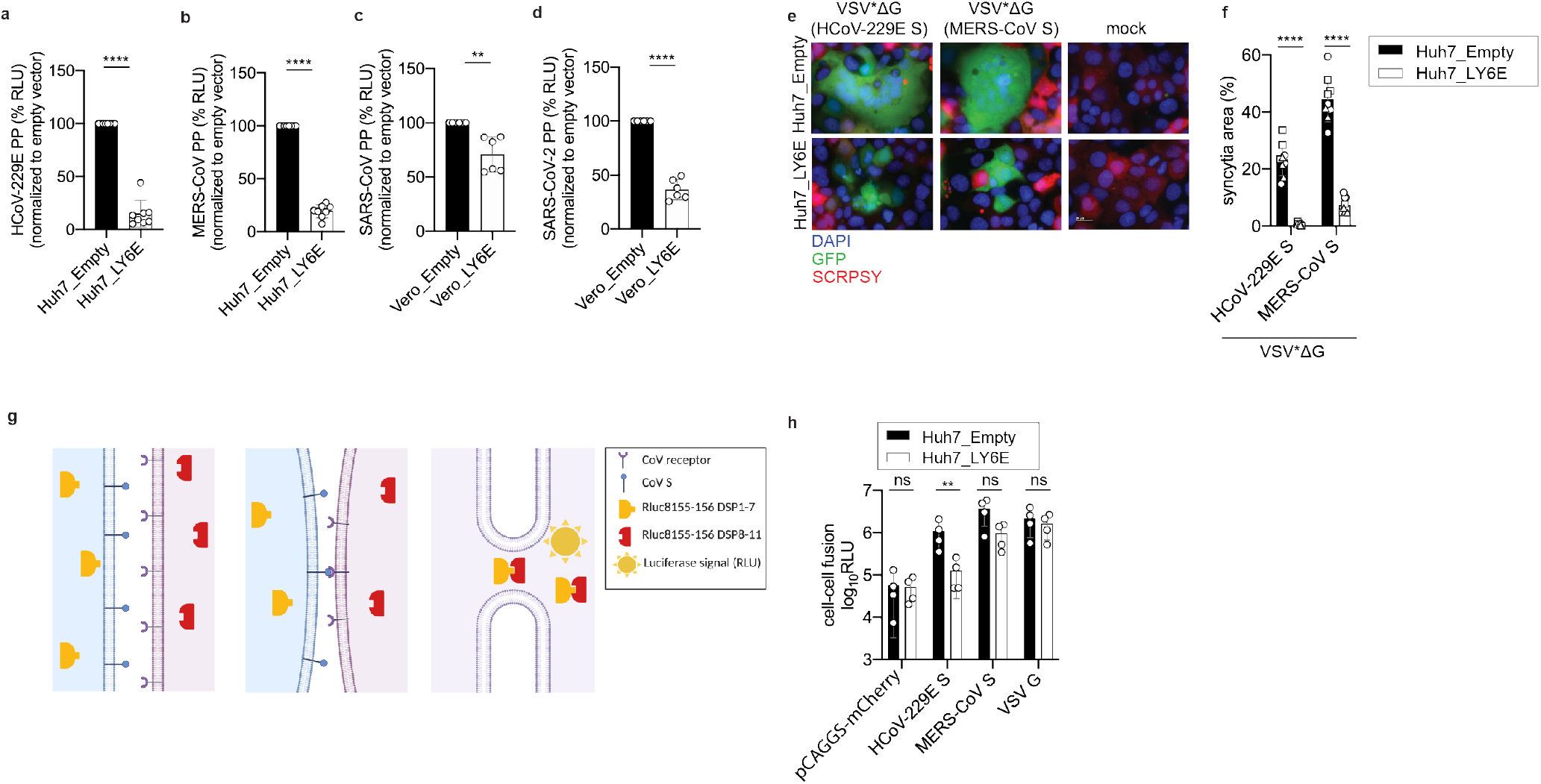
LY6E inhibits viral membrane fusion and syncytia formation. **a-d**, VSV pseudoparticles (PP) harboring spike proteins from HCoV-229E (**a**), MERS-CoV (**b**), SARS-CoV (**c**), SARS-CoV-2 (**d**) were inoculated on LY6E- or empty vector-expressing cells. Virus entry efficiency was quantified by measuring virus-encoded luciferase reporter activity. **e**, VSV pseudoparticles expressing VSV G protein on their surface and encoding CoV S protein and GFP (VSV*ΔG(CoV S)) were inoculated with LY6E- or empty vector-expressing Huh7. Syncytia formation was analyzed by immunofluorescence microscopy. Blue: DAPI, green: GFP, red: TagRFP inserted in SCRPSY vector. **f**, Quantification of VSV*ΔG(CoV S) induced syncytia depicted as percentage syncytia area. Three independent areas were analyzed per biological replicate (circle, square, triangle). **g**, Schematic depiction of the cell-cell fusion assay created with BioRender. **h**, Cell-cell fusion was measured by co-incubating cells transfected with plasmids encoding CoV S proteins and a split-luciferase construct (DSP1-7) together with CoV permissive LY6E-expressing or control cells transfected with the second fragment of split-luciferase (DSP8-11). Data represent averages of independent biological replicates, n=8 (**a,b**), n=6 (**c,d**), n=3 (**f**), n=4 (**h**). Statistical significance was determined by unpaired student’s t-test with Welch’s correction (**a-d**) and 2way ANOVA followed by Sidak’s multiple comparison test (**f**,**h**). Outliers were removed using Grubbs outlier test (**a**). In **e**, scale bar is 20 µM. Error bars: SD. *P* values: **a**, **** p=<0.0001; **b**, **** p=<0.0001; **c**, **p=0.0066; **d**, **** p=<0.0001; **f**, **** p=<0.0001; **h**, ns p=0.9975, ** p=0.0054, ns p=0.1520, 0.9892.

Next, we aimed to evaluate LY6E-mediated CoV restriction *in vivo*. Systemic genetic ablation of *Ly6e* in mice is embryonic lethal^18^. Therefore, we generated transgenic *Ly6e^fl/fl^* mice, which are functionally ‘wild-type (WT)’ for Ly6e expression, and crossed them with *Vav1-iCre* mice to ablate *Ly6e* in cells that originate from a hematopoietic stem cell (HSC) progenitor (**Extended Data Fig. 5a-b**). This genetic model was chosen since immune cells are critical for protection from CoV *in vivo*^5,19–23^. Bone-marrow derived macrophages (BMDM) from *Ly6e^ΔHSC^* mice had reduced *Ly6e* mRNA levels (**Extended Data Fig. 5c**). The natural mouse pathogen MHV is a well-studied model CoV that causes hepatitis and encephalomyelitis in mice^24^. *Ly6e^ΔHSC^* BMDM were more susceptible to MHV infection than *Ly6e^fl/fl^* BMDM, demonstrating that murine Ly6e is a restriction factor for murine CoV (**Extended Data Fig. 5d**).

To determine whether Ly6e is important for controlling CoV infection *in vivo*, *Ly6e^fl/fl^* and *Ly6e^ΔHSC^*mice were infected with a high dose (5 × 10^4^ PFU) of MHV that causes sub-lethal hepatitis in WT C57BL/6J mice^25^. Since MHV pathogenesis is influenced by sex^26^, we performed independent experiments in male and female mice. *Ly6e^ΔHSC^* mice of both sexes rapidly succumbed to MHV infection by day 6 (**Fig. 3a-b**, **Extended Data Fig. 6a-b**). To examine viral pathogenesis and immunopathology in infected mice, *Ly6e^fl/fl^* and *Ly6e^ΔHSC^* mice were infected with the same high dose of MHV and euthanized 3- or 5-days post-infection. At both time points, female *Ly6e^ΔHSC^* mice exhibited significantly higher levels of liver damage as measured by serum ALT while no difference was observed in male mice (**Fig. 3c,i**, **Extended Data Fig. 6c,i**). However, hepatic viral burden did not differ in either sex (**Fig. 3d,j**, **Extended Data Fig. 6d,j**), possibly due to tissue saturation at this high dose. Indeed, female but not male *Ly6e^ΔHSC^* mice exhibited increased liver damage and viral burden at a low dose (5 PFU) of MHV (**Extended Data Fig. 7a-b, f-g**). Spleen viral burden was elevated in female and male *Ly6e^ΔHSC^* mice at both time points and doses of MHV (**Fig. 3e,k**, **Extended Data Fig. 6e,k**, **Extended Data Fig. 7c,h**). At 3 days post-infection, there was no difference in liver necrosis, but inflammation was moderately reduced in female but not male *Ly6e^ΔHSC^* mice (**Fig. 3f-h**, **Extended Data Fig. 6f-h**). By 5 days post-infection and at both low and high doses of MHV, female *Ly6e^ΔHSC^* mice had significantly higher levels of liver necrosis and a reduced presence of mononuclear immune cells (**Fig. 3l-n**, **Extended Data Fig. 7d-e**), whereas no difference was observed in male mice (**Extended Data Fig. 6l-n**, **Extended Data Fig. 7i-j**). Liver damage in *Ly6e^ΔHSC^* mice was further corroborated at an intermediate viral dose (5 × 10^3^ PFU), which resulted in visibly apparent brittleness, pallor, and necrotic foci (**Extended Data Fig. 7k-n**). These data demonstrate that Ly6e in hematopoietic cells is important for controlling murine CoV infection.

**Figure 3.**
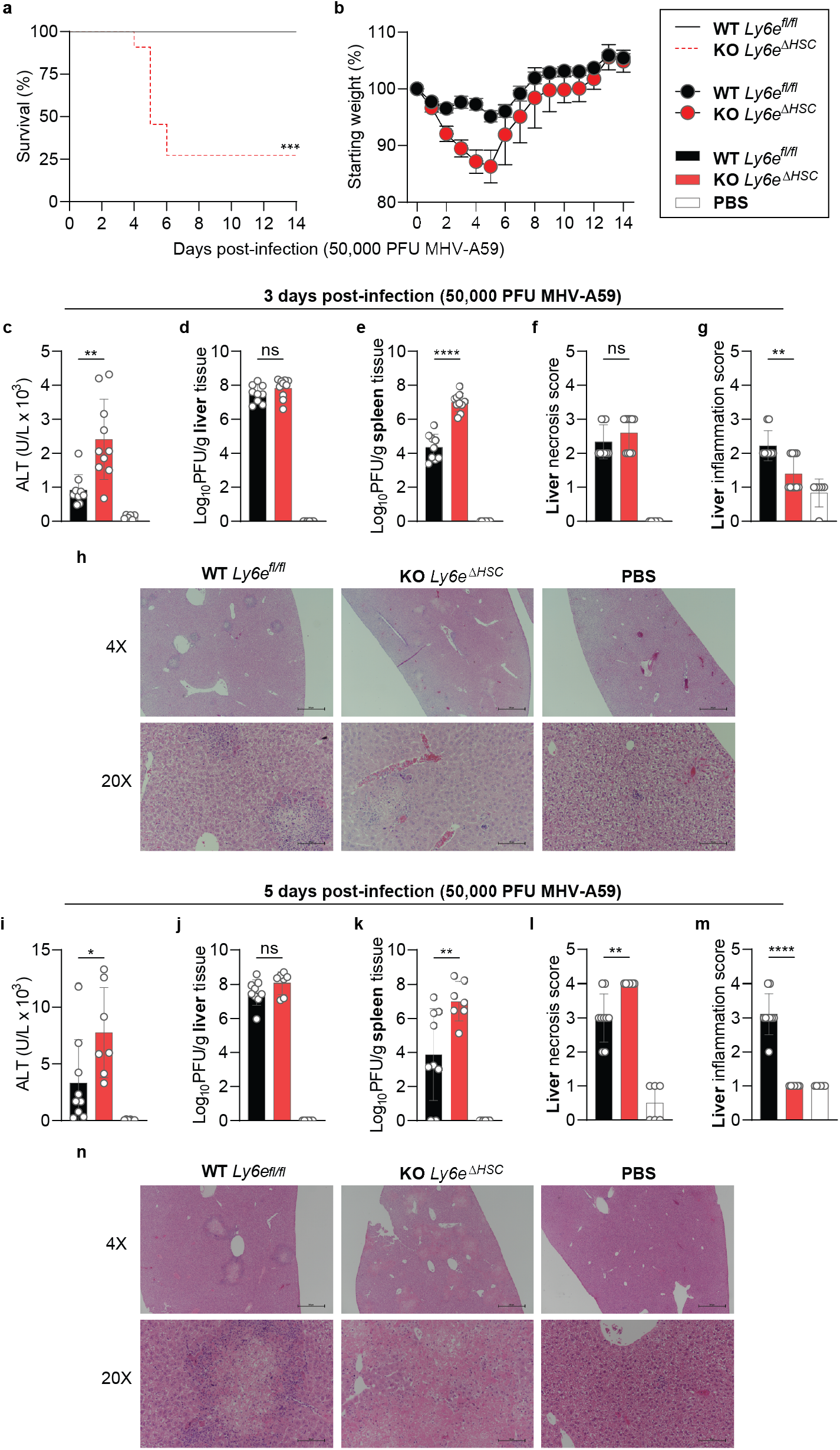
*Ly6e^ΔHSC^* mice are more susceptible to murine hepatitis virus. **a-b,** Female *Ly6e^fl/fl^* and *Ly6e^ΔHSC^* mice were injected with PBS or MHV and monitored for survival (**a**) and weight loss (**b**). **c-n**, Mice were assessed at 3 (**c-h**) or 5 (**i-n**) days post-infection for serum ALT (**c,i**), viral titers in liver (**d,j**) and spleen (**e,k**), and liver necrosis and inflammation (**f-h**, **l-n**). For **a-b**, n=13 for days 0-14 (*Ly6e^fl/fl^*), n=11 for day 0-3, n= 10 for days 4-5, n=5 for days 5-6, n=3 for days 6-14 (*Ly6e^ΔHSC^*) from three pooled experiments. For **c-g**, n=9 (*Ly6e^fl/fl^*), n=10 (*Ly6e^ΔHSC^*), n=6 (PBS) from two pooled experiments. For **i-m**, n=9 (*Ly6e^fl/fl^*,), n=7 (*Ly6e^ΔHSC^*), n=6 (PBS) from two pooled experiments. In **h,n**, scale bars are 500 µM (4x) and 100 µM (20x). Statistical significance was determined by Mantel-Cox test (**a**), two-tailed unpaired student’s t-test with Welch’s correction (**c-e**, **i-k**), two-tailed Mann-Whitney U test (**f-g**, **l-m**). Error bars: SEM (**b**), SD (**c-g**, **i-m**). *P* values: **a**, *** p=0.0002; c, ** p=0.0031; **d**, ns p=0.2136; **e**, **** p=5.759 × 10^−7^; **f**, ns p=0.3698; **g**, ** p=0.0054; **i**, * p=0.0430; **j**, ns p=0.1321; **k**, ** p=0.0094; **l**, ** p=0.0072; **m**, **** p=8.741 × 10^−5^.

We next used a transcriptomic approach to evaluate the effect of *Ly6e* ablation on global gene expression in liver and spleen of female *Ly6e^fl/fl^* and *Ly6e^ΔHSC^* mice injected with PBS or the high dose of MHV at 3- and 5-days post-infection. In infected tissues, loss of *Ly6e* in hematopoietic cells correlated with differential gene expression in pathways associated with tissue damage, such as liver-specific metabolic genes, angiogenesis, wound healing, and immune response to viruses (**Fig. 4a-c**). We also observed a striking loss of genes associated with the type I IFN response, inflammation, antigen presentation, and B cells in infected *Ly6e^ΔHSC^* mice (**Fig. 4d-e**, **Extended Data Fig. 8**).

**Figure 4.**
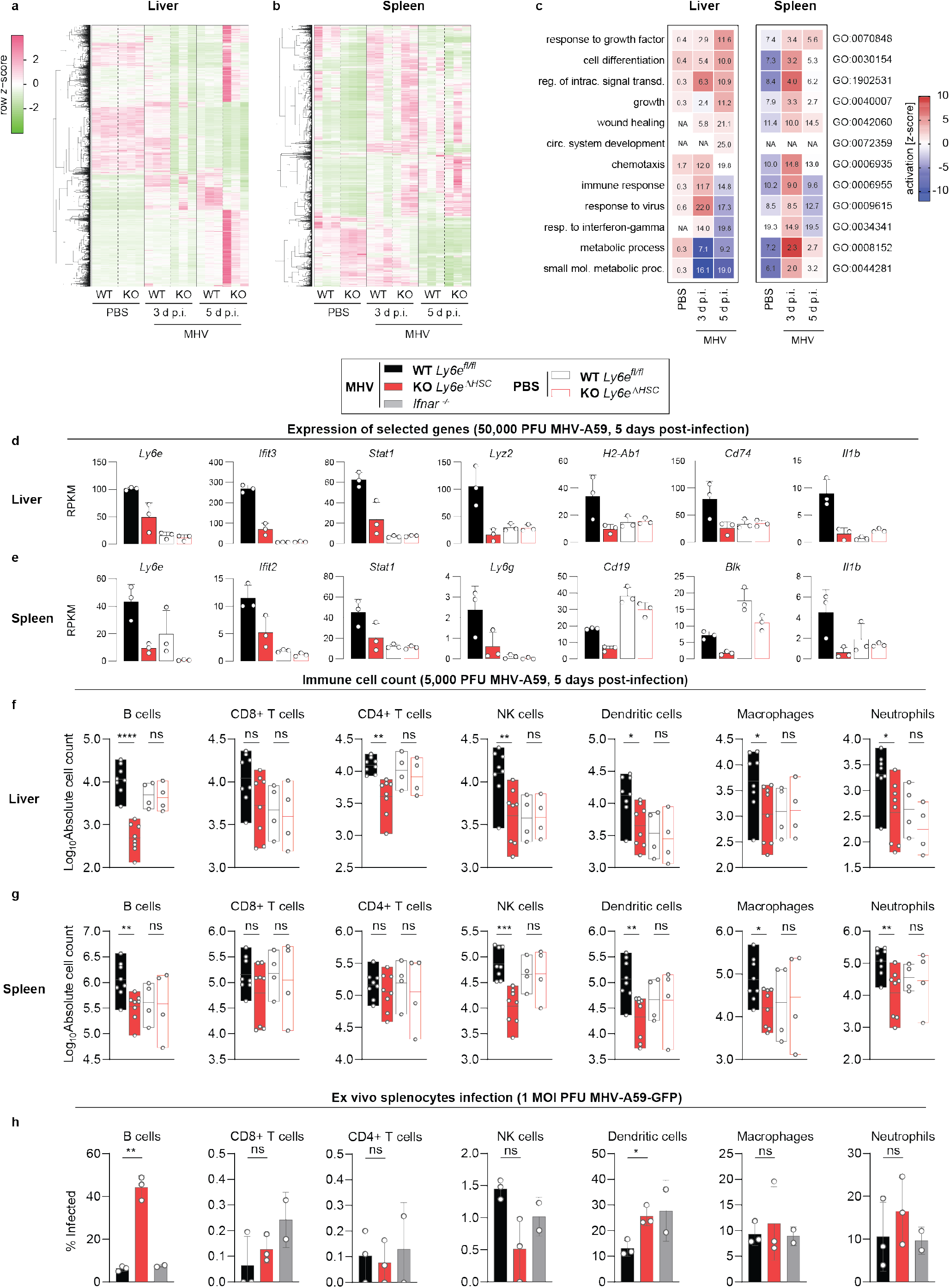
Murine hepatitis virus infection alters hepatic and splenic immune responses in female *Ly6e^ΔHSC^* mice. **a-e**, Transcriptomic analysis from *Ly6e^fl/fl^* (WT) and *Ly6e^ΔHSC^* (KO) mice. **a-b**, Heat maps displaying significant changes (mean RPKM > 0.5; fold change > 2; FDR ≤ 0.05) in liver (**a**) and spleen (**b**). Dendrograms are normalized read data (row z-score) clustered with complete linkage method employing Spearman Rank correlation distance measurement. **c**, Pathway analysis of KO versus WT. Up-(red) or down-regulated (blue) pathways indicated by activation z-score. Numbers show significantly dysregulated genes as percentage of total gene number included in pathway. **d-e**, Expression of select genes in liver (**d**) and spleen (**e**). **f-g**, Immune cell counts from liver (**f**) and spleen (**g**). **h**, Infection of cultured splenocytes. For **a-e**, n=3. For **f-g**, n=8 MHV-injected, n=4 PBS-injected from two pooled experiments. For **h**, n=3 (*Ly6e^fl/fl^* and *Ly6e^ΔHSC^*) or n=2 (*Ifnar^−/−^*) from two pooled experiments. Significance for **f-h** was determined by two-tailed unpaired student’s t-test with Welch’s correction. Error bars: SD (**d-h**). *P* values (left-right): **f**, **** p=1.3644 × 10^−6^, ns p=0.7952, ns p=0.0781, ns p=0.7732, ** p=0.0016, ns p=0.6261, ** p=0.0068, ns p=0.9494, * p=0.0141, ns p=0.7639, * p=0.0436, ns p=0.9600, * p=0.0174, ns p=0.3096; **g,** ** p=0.0052, ns p=0.9447, ns p=0.1579, ns p=0.7769, ns p=0.4104, ns p=0.6924, *** p=1.754 × 10^−4^, ns p=0.9842, ** p=0.0034, ns p=0.9848, * p=0.0169, ns p=0.8591, ** p=0.0076, ns p=0.8623; **h,** ** p=0.0051, ns p=0.4525, ns p=0.7379, ns p=0.0702, * p=0.0119, ns p=0.6787, ns p=0.4112.

The histopathology (**Fig. 3h,n**) and transcriptome data (**Fig 4a-e**, **Extended Data Fig. 8**) prompted us to determine how loss of *Ly6e* affects immune cell numbers during infection. Absolute numbers of different immune cell subsets were quantified from *Ly6e^ΔHSC^* and *Ly6e^fl/fl^* mice 5 days after injection with PBS or the intermediate dose of MHV (**Extended Data Fig. 7k-n**, **Extended Data Fig. 9**). Multiple cell types were depleted in livers and spleens of infected *Ly6e^ΔHSC^* mice, irrespective of sex (**Figure 4f-g, Extended Data Fig. 10a-b**). B cells were dramatically reduced in livers from infected *Ly6e^ΔHSC^* mice. Hepatic CD4^+^ T cells, NK cells, dendritic cells (DC), macrophages, and neutrophils were also depleted in infected *Ly6e^ΔHSC^* mice, whereas CD8^+^ T cells were unchanged (**Fig. 4f**, **Extended Data Fig. 10a**). Infected *Ly6e^ΔHSC^* mice had a reduction in all splenic immune cell subsets except for CD4^+^ and CD8^+^ T cells (**Fig. 4g**, **Extended Data Fig. 10b**). Depletion of immune cell populations in *Ly6e^ΔHSC^* organs was MHV-dependent, as cell numbers were not altered in PBS-injected mice.

To determine whether loss of immune cells in MHV-infected *Ly6e^ΔHSC^* mice correlates with increased permissiveness to infection, we assessed MHV infection in splenocytes cultured from *Ly6e^ΔHSC^, Ly6e^fl/fl^*, and *Ifnar^−/−^* mice. In *Ly6e^fl/fl^* splenocyte cultures, MHV infected macrophages, neutrophils, DC, and B cells, but not CD4^+^ or CD8^+^ T cells, as previously published (**Fig. 4h**, **Extended Data Fig. 10c**)^23^. Strikingly, B cells from both male and female *Ly6e^ΔHSC^* mice were highly susceptible to MHV infection when compared to B cells from *Ly6e ^fl/fl^* or *Ifnar^−/−^* mice. DC from female *Ly6e^ΔHSC^* mice were also more susceptible to MHV infection. Surprisingly, no other cell type demonstrated a strong Ly6e-dependent susceptibility to MHV infection.

Together, our data demonstrate an important role for Ly6e in immune cell-mediated control of CoV infection. Higher viral burden and lower liver inflammation suggests that liver damage in MHV-infected female *Ly6e^ΔHSC^* mice may be due to viral pathogenesis that exceeds the ability of the immune system to control the infection. Notably, we observed MHV-induced loss of three classes of cells: 1) cells that are not permissive to the virus (CD4^+^ T cells), 2) cells that are permissive to MHV but do not demonstrate Ly6e KO-dependent infection (NK cells, macrophages, and neutrophils), and 3) cells that are permissive to MHV in a Ly6e KO-dependent manner (B cells and DC). This latter group is capable of antigen-presentation to CD4^+^ T cells^27,28^. Thus, in the absence of Ly6e, unrestricted MHV infection of B cells or DC may impair T cell priming, thereby negatively impacting recruitment and activation of NK cells, macrophages, and neutrophils that contribute to control of viral disease.

In conclusion, we identified LY6E as a pan CoV restriction factor that limits CoV entry and protects the host from severe viral disease. Our findings are striking given that LY6E has been primarily associated with an enhancing phenotype in which LY6E promotes entry of multiple viruses^29^. However, our data clearly expand the role of LY6E during coronaviral infection and establishes a novel function in protecting the host immune cell compartment. Determining the precise molecular mechanism underlying how LY6E inhibits CoV S protein-mediated membrane fusion will advance our understanding of cellular antiviral defenses against these important human pathogens. Antiviral membrane fusion inhibitors have been successfully implemented for treatment of HIV-1 infection^30^. A therapeutic approach mimicking the mechanism of action of LY6E could provide a first line of defense against novel emerging CoV infections. Furthermore, delineating which Ly6e-expressing immune cells protect mice from MHV will provide new insight into how individual antiviral effectors in distinct cellular compartments modulate viral pathogenesis.

## Methods

### Viruses

Recombinant HCoV-229E^31^, HCoV-229E-Rluc (expressing Renilla luciferase [Rluc] by replacing the majority of HCoV-229E ORF4)^32^, HCoV-OC43^33^, MHV strain A59^34^, MHV-Gluc, strain A59 (expresses a Gaussia luciferase [Gluc] replacing accessory gene 4)^35^, MHV-GFP strain A59^25^, MERS-CoV strain EMC^36^, and SARS-CoV strain Frankfurt-1^37^ have been described previously. SARS-CoV-2 (BetaCoV/Wuhan/IVDC-HB-01/2019, GISAID Accession ID: EPI_ISL_402119) stocks were propagated on VeroE6 cells. HCoV-229E viruses were propagated on Huh-7 cells, MERS-CoV and SARS-CoV were propagated on VeroB4 cells, HCoV-OC43 was propagated on HCT-8 cells^38^, and MHV stocks were propagated on 17Cl1 cells. Lentivirus particles using the SCRPSY lentiviral backbone were generated as described previously^7,8^.

VSV*ΔG(Fluc) (G glycoprotein-deficient VSV encoding green fluorescent protein [GFP] and firefly luciferase [Fluc]) was generated as described previously and was propagated on BHK-G43 cells^39^. The generation of viral stocks for the following viruses has been previously described: hPIV-3-GFP^40^ (based on strain JS, generously provided by P.L. Collins), RSV-GFP^41^ (based on strain A2, generously provided by P.L. Collins), YFV-Venus^42^ (derived from YF17D-5′C25Venus2AUbi), DENV-GFP^43^ (derived from IC30P-A, a full-length infectious clone of strain 16681), WNV-GFP^44^ (derived from pBELO-WNV-GFP-RZ, generously provided by I. Frolov), HCV-Ypet^45^ (based on the chimeric Jc1 virus of strains: J6 and JFH-1), SINV-GFP^46^ (derived from pS300/pS300-GFP, generously provided by M.T. Heise), VEEV-GFP^47^ (derived from pTC83-GFP infectious clone, generously provided by I. Frolov), CHIKV-GFP^48^ (derived from pCHIKV-LR 5′GFP, generously provided by S. Higgs). ZIKV (PRVABC59, obtained from the CDC, Ft. Collins) was amplified and titrated as described previously^49^.

### Viral infection assays

HCoV-229E: 1.5 × 10^5^ stable LY6E expressing or control cells were seeded in a 12-well plate and infected with HCoV-229E in a serial dilution. Cells were incubated for 2 hours at 33°C, then viral supernatant was removed, and an overlay of 1:1 Avicel (2.4%) (Avicel RC-581NF) and 2x DMEM, supplemented with 10% FBS and 1x penicillin/streptomycin (p-s), was added. Cells were incubated for 3 days at 33°C before medium was removed, cells were washed, and stained with crystal violet (Sigma-Aldrich). Plaque forming units (PFU) were calculated.

HCoV-OC43: 5.0 × 10^4^ stable LY6E expressing or control cells were seeded in a 24-well plate and infected at 33°C for 1 hour with HCoV-OC43 (MOI=1). Virus was aspirated and 10% FBS/1x NEAA/1x p-s/RPMI (cRPMI) was added back to cells. 24 hours post-infection, cells were dissociated using Accumax (Sigma-Aldrich), fixed in 1% PFA, permeabilized per manufacturer’s protocol (BD Cytofix/Cytoperm), and stained for nucleoprotein (1:500) and a goat anti-mouse AlexaFluor488-conjugated secondary antibody (1:2000). Infection was analyzed by flow cytometry.

MERS-CoV/ SARS-CoV/ SARS-CoV-2: 1 × 10^4^ stable LY6E expressing or control cells were seeded in a 96-well plate and infected at 37°C in a serial dilution assay. 3-4 days post-infection supernatant was removed, and cells were stained with crystal violet (Sigma-Aldrich). Cytopathic effect (CPE) was determined upon visual inspection and TCID_50_/ml calculated according to the Reed and Muench method. MHV-Gluc: 1 × 10^5^ stable LY6E expressing or control cells were seeded in a 24-well plate and infected with MHV-Gluc (MOI=0.1) at 37°C. After 2 hours, virus inoculum was removed, cells were washed with PBS, and cRPMI added back. 24 hours post-infection, cell culture supernatant was harvested, and Gluc activity was measured using Pierce Gaussia Luciferase Glow Assay Kit (ThermoFisher Scientific) and a plate luminometer (EnSpire 2300 Multilabel reader by Perkin Elmer). For testing the virus panel against LY6E, 1 × 10^4^ stable LY6E expressing or control Huh7.5 cells were seeded in a 96-well plate. Cells were infected with the following viruses and harvested after the indicated time: HCoV-229E (MOI=0.001, 72 hours), HCV-Ypet (MOI=1, 72 hours), CHIKV-GFP (MOI=0.005, 24 hours), hPIV-3-GFP (MOI=0.015, 24 hours), RSV-GFP (MOI=1.5, 24 hours), SINV-GFP (MOI=0.1, 24 hours), VEEV-GFP (MOI=0.0075, 24 hours), WNV-GFP (MOI=1.5, 24 hours), ZIKV PRVABC59 (MOI=0.01, 48 hours), YFV 17D-venus (MOI=0.01, 48 hours) and DENV-GFP (MOI=0.05, 72 hours). At the indicated time points, cells were fixed with 4% paraformaldehyde (PFA)/PBS. ZIKV-infected cells were stained for viral E protein and an AlexaFluor488-conjugated goat anti-mouse secondary antibody. Images were acquired with a fluorescence microscope and analyzed using ImageXpress Micro XLS (Molecular Devices, Sunnyvale, CA).

### HCoV-229E-Rluc infection assay of KO cells

For infection assays, 1 × 10^5^ target cells were seeded in a 24-well plate one day prior infection. Cells were infected with HCoV-229E-Rluc (MOI 0.1) in OptiMEM (Gibco) for 2 hours at 33°C. Cells were washed 1x with PBS and 10% FBS/1x NEAA/1x p-s/DMEM (cDMEM) was added back. Cells were incubated at 33°C for 24 hours, then washed with PBS and lysed using the Renilla Luciferase Assay System kit (Promega). Rluc activity was measured using a plate luminometer (EnSpire 2300 Multilabel reader by Perkin Elmer). For the reconstitution with CRISPR resistant LY6E (CR LY6E), 2.5 × 10^4^ cells were seeded in a 24-well plate. One day post-seeding, cells were either left untransduced or transduced with a lentiviral vector encoding for CR LY6E or the empty control. 48 hours post-transduction, cell lysates were either infected with HCoV-229E-Rluc (MOI=0.1) for 24 hours as described above or harvested for Western blot.

### VSV pseudoparticles

Pseudotyping of VSV*ΔG(Fluc) was performed as previously described^50^. Briefly, 6 × 10^5^ 293LTV cells were seeded in a 6-well plate and transfected using Lipofectamine 2000 (Invitrogen) which was complexed with DNA plasmids driving the expression of either VSV G protein (positive control), the respective CoV S proteins, or the fluorophore mCherry (negative control). Expression vectors for vesicular stomatitis virus (VSV, serotype Indiana) glycoprotein (VSV-G, Genbank accession number: NC_001560), HCoV-229E S (pCAGGS-229E S, Genbank accession number: X16816), MERS-CoV S (pCAGGS-MERS S, Genbank accession number: JX869059 with silent point mutation (C4035A, removing internal XhoI), SARS-CoV S (pCAGGS-SARS S, Genbank accession number: AY291315.1 with two silent mutation (T2568G, T3327C)), have been described previously^51–53^. The expression plasmid for SARS-CoV-2 was generated as follows: the coding sequence of a synthetic, codon-optimized (for human cells) SARS-CoV-2 DNA (GeneArt Gene Synthesis, Thermo Fisher Scientific) based on the publicly available protein sequence in the National Center for Biotechnology Information database (NCBI Reference Sequence: YP_009724390.1) was PCR-amplified and cloned into the pCG1 expression vector between BamHI and XbaI restriction sites. The MERS-CoV S cleavage mutants have been described before^17^. pCAGGS_mCherry was generated upon cloning of the mCherry reporter gene into the pCAGGS expression vector (primer sequences and cloning procedure are available upon request). At 20 hours post-transfection, cells were infected with VSV-G-trans-complemented VSV*ΔG(FLuc) (MOI=5) for 30 minutes at 37°C, washed with PBS, and further incubated for 1 hour in cDMEM that was supplemented with Mab I1 (ATCC), a neutralizing monoclonal antibody directed to the VSV G protein. Cells were then washed and cDMEM was added back. Viral supernatant was harvested at 16 – 24 hours post-infection. Cellular debris was removed by centrifugation (3,000 × *g* for 10 minutes) and used to inoculate 2 × 10^4^ target cells in a 96-well plate (6 technical replicates). Cells were incubated for 16–18 hours at 37°C before Fluc activity in cell lysates was detected using a plate luminometer (EnSpire 2300 Multilabel reader by Perkin Elmer) and the Bright-Glo Luciferase Assays System (Promega). Experiments with pseudotyped SARS-CoV and SARS-CoV-2 were performed as previously described^17^.

### Split luciferase fusion assay

The split luciferase fusion assay has been described before, with slight modifications^54^. Briefly, 6 × 10^5^ 293LTV cells were transfected with plasmids encoding CoV S proteins (pCAGGS-HCoV-229E, pCAGGS-MERS-CoV) together with a split-luciferase construct (Rluc8155-156 DSP1-7). The empty expression plasmid (pCAGGS-mCherry) as well as a construct encoding for VSV G served as negative and positive control, respectively. 2 × 10^5^ stable LY6E expressing or empty control Huh7 cells were transfected using Lipofectamine 2000 (Thermo Fisher Scientific) with a plasmid encoding the second half of the split-luciferase protein (Rluc8155-156 DSP8-11). Approximately 30 hours post-transfection, both cell populations were dissociated (TryPLE Express, Gibco), counted, and equal cell numbers (2×10^4^ cells each) were co-cultured for 16 – 20 hours at 37°C. Supernatant was harvested and cells lysed using ice-cold passive lysis buffer (Promega). Cells were immediately transferred to −80°C for at least 1 hour, before Rluc activity was determined using the Renilla Luciferase Assay System (Promega). Cells were kept on ice during all times post-cell lysis to eliminate DSP post-lysis complementation.

### Generation of recombinant VSV vector driving CoV S protein expression (VSV*ΔG(CoV))

Following extraction of total RNA from MERS-CoV (strain EMC) infected VeroE6 cells, the cDNA encoding the MERS-CoV S protein was generated by reverse transcription (RevertAid Premium Reverse Transkriptase, ThermoScientific). Three overlapping cDNA fragments were amplified by PCR (Phusion Hot Start II High Fidelity Polymerase, Thermo Scientific) and subsequently assembled by overlapping PCR. The cDNAs encoding the S proteins of either MERS-CoV (strain EMC) or HCoV-229E (truncated variant lacking the 10 amino acids at the C terminus, original plasmid pCAGGS-229E S) were amplified by PCR and inserted into the pVSV* plasmid^55^ between MluI and BstEII restriction sites. The resulting plasmids were used to generate the recombinant viruses VSV*ΔG(MERS S) and VSV*ΔG(229E S) according to a published procedure^56^. All viruses were propagated on BHK-G43 cells^57^, resulting in viruses predominantly containing the homotypic VSV glycoprotein G in the envelope.

### Syncytia formation assay

Cells expressing LY6E or empty control were infected with VSV*ΔG(MERS S) or VSV*ΔG(229E S) at a MOI of 0.01. At 20 hours post-infection, the cells were fixed with 3% PFA/PBS and the nuclei stained with 4’,6-diamidine-2’-phenylindole (DAPI, Sigma). An inverted fluorescence microscope (Zeiss) was used to determine area percentage area covered by syncytia.

For heterologous cell-cell fusion assay, BHK-21 cells were infected with VSV*ΔG(CoV S) at a MOI of 1. After 2 hours, the cells were treated with trypsin/EDTA (Life Technologies, Zug, Switzerland) and resuspended in DMEM supplemented with 5% FBS (2×10^4^ cells/mL). The infected BHK-21 cell suspension (500 µL) was seeded into 24-well cell culture plates along with LY6E- or empty control-expressing Huh7 (1 × 10^4^ cells). Cell co-cultures were incubated for 7 hours at 37°C, fixed with 3.6% formaldehyde diluted in PBS, (Grogg-Chemie AG, Stettlen, Switzerland) and stained with DAPI. The percentage area covered by syncytia was calculated as above.

### Mice

Ly6e^tm1a^ ES cells were obtained from the EUCOMM consortium^58^ and microinjected into C57BL/6J blastocysts by the UTSW Transgenic Technology Center. Chimeric mice with germline transmission were bred to obtain *Ly6e^tm1a/+^* offspring. *Ly6e^tm1a/+^* mice were crossed with FLPe-expressing mice (*B6N.129S4-Gt(ROSA)26Sor^tm1(FLP1)Dyn^/J, #*16226, Jackson Laboratories) to obtain *Ly6e^fl/+^* offspring, which were bred to homozygosity. Conditional *Ly6e^fl/fl^* mice were bred to *Vav1-iCre* transgenic mice (B6.Cg-Commd10^Tg(Vav1-icre)A2Kio^/J, #008610, Jackson Laboratories) to obtain *Ly6e^ΔHSC/+^* (*Ly6e^fl/+^; Vav1-iCre*) offspring. *Ly6e^ΔHSC/+^* mice were bred to obtain *Ly6e^ΔHSC^* (*Ly6e^fl/fl^; Vav1-iCre*) offspring, which harbor a deletion of exon 3 and 4 in hematopoietic stem cells (HSC). Experimental animals were obtained by crossing *Ly6e^ΔHSC^* and *Ly6e^fl/fl^* mice. Genotype was confirmed by PCR of genomic DNA in-house or outsourced to Transnetyx. Ablation of *Ly6e* was confirmed by qPCR in immune cells from spleen, liver, or bone marrow. Animal studies were carried out in specific-pathogen-free barrier facilities managed and maintained by the UTSW Animal Resource Center. All procedures used in this study complied with federal and institutional guidelines enforced by the UTSW Institutional Animal Care and Use Committee (IACUC).

### *In vivo* infection, viral titers, liver ALT, and liver histology

Six to twelve-week-old female and male mice were injected intraperitoneally with MHV-A59 diluted in PBS to the indicated titers or PBS for a mock infection control. All infected mice were monitored daily for weight and mortality. Animals that lost more than 20% of their original body weight were euthanized per IACUC guidelines.

Viral titers in liver and spleen were determined from frozen organs after weighing, homogenization, and plaque assay on L929 cells. Alanine aminotransferase (ALT) was measured in fresh, unfrozen serum using VITROS MicroSlide Technology by the UTSW Mouse Metabolic Core. Livers were fixed in 10% neutral buffer formalin, embedded in paraffin, sectioned at 5 µM, and stained with hematoxylin and eosin (H&E). Slides were analyzed by an independent pathologist (UTSW Animal Resource Center) who was blinded to experimental conditions. A numerical score was assigned for degree of inflammation and degree of necrosis for each liver. Inflammatory cell infiltration was scored using the following criteria: 0 = none, 1 = minimal, 2 = mild, 3 = moderate, and 4 = marked. The necrosis score was defined by the percent necrosis. 0 = none, 1 = <2%, 2 = 2-20%, 3 = 21-40%, and 4 = >40%.

### Primary cell preparation for flow cytometry

All primary cells were maintained in 10% FBS/10 mM HEPES/1 mM sodium pyruvate/2 mM L-glutamine/1x p-s/RPMI (cRPMI2) unless otherwise indicated.

Primary bone marrow derived macrophages (BMDM) were prepared as described previously^60^. Viable cells were quantified using trypan blue exclusion. BMDM were plated at 2.5 × 10^4^ cells per well in non-treated 24-well tissue culture plates. Cells were infected the next day with 0.1 MOI MHV-A59-GFP and isolated 6 hours post-infection for analysis by flow cytometry.

Splenocytes were prepared by mashing whole spleens against a 70 µM cell strainer. Red blood cells were removed by using RBC lysis buffer (Tonbo Biosciences). Viable cells were quantified using trypan blue exclusion. For *in vitro* infection, splenocytes were plated at 1×10^6^ cells per well in non-treated 6-well tissue culture plates. Cells were infected the next day with 1 MOI MHV-A59-GFP and isolated 6 hours post-infection. Infected cells were stained with a fluorescent viability dye (Ghostdye Violet 450, Tonbo Biosciences) and fixed in 1% PFA/PBS. The next day, fixed cells were treated with anti-CD16/CD32 (Tonbo Biosciences) and then stained with antibodies that recognize surface lineage markers and analyzed by flow cytometry.

Livers were perfused by PBS injection into the inferior vena cava and out of the portal vein. Blanched livers were minced with sterile scissors and mashed against a 70 µM cell strainer. Hepatocytes and large debris were pelleted at 60 × *g* for 1 minute, 22°C, with no brake. Intrahepatic immune cell (IHIC) and residual hepatocyte containing supernatants were pelleted at 480 × *g* for 8 minutes, 22°C, with max brake and resuspended in 37.5% Percoll (Sigma-Aldrich)/cRPMI2. Remaining hepatocytes were separated from IHIC by centrifugation at 850 × *g* for 30 minutes, 22°C, with no brake. Red blood cells were removed from the resulting pellet with RBC lysis buffer (Tonbo Biosciences). Viable cells were quantified using trypan blue exclusion. IHIC and splenocytes (4 × 10^5^) from infected or mock-infected mice were stained with a fluorescent viability dye (Ghostdye Violet 450, Tonbo Biosciences), treated with anti-CD16/CD32 (Tonbo Biosciences), stained with antibodies that recognize surface lineage markers, and then fixed in 1% PFA/PBS. Fixed volumes of cell suspensions were analyzed by flow cytometry the next day. Absolute cell counts were determined by multiplying final number of lineage marker-positive cells by dilution factor used to normalize cell counts for antibody staining.

All primary cell samples were resuspended in 3% FBS/PBS and analyzed using a S1000 Flow Cytometer with a A600 96-well plate high throughput extension and compensated using CellCapture software (Stratedigm). Data was analyzed with FlowJo Software (Treestar). The flow gating strategy for liver and spleen immune cells is included in **Extended Data Figure 9**.

### Statistical analysis

Differences in data were tested for significance using GraphPad Prism v8.3.1 for Windows (GraphPad). For details regarding the statistical tests applied, please refer to the figure legends. *P* values < 0.05 were considered significant.

### Data availability

The authors declare that the data supporting the findings of this study are available within the article and its Supplementary Information files or are available on request. The RNAseq data discussed in this publication have been deposited in the Gene Expression Omnibus (GEO) database, https://www.ncbi.nlm.nih.gov/geo (GSE146074).

## Acknowledgements

We would like to thank the following people for their generous contribution of reagents and infrastructure which greatly enhanced this study, especially the sequencing facility in Bern, the diagnostic facility of IVI, and the Animal Resource Center at UTSW. Furthermore, we are grateful to Robin Pohorelsky, Shay Elias, and Nina Garcia from the UTSW Animal Resource Center for assisting in the *in vivo* experiments. We are grateful to Samira Locher (IVI, Bern, Switzerland) for cloning the MERS spike cDNA. Furthermore, we would like to thank Berend Jan Bosch, Marcel Müller and Christian Drosten for reagents and the SARS-CoV-2 virus stocks. We thank the following investigators for contributing viral molecular clones or viral stocks: P.L. Collins (RSV, hPIV-3), I. Frolov (VEEV), M.T. Heise (SINV), S. Higgs (CHIKV), and the CDC (ZIKV).

Finally, we’d like to thank all members, past and present, of Institute of Virology and Immunology, the Schoggins lab, and the Rice lab for useful discussions.

S.P. was supported by the European Commission’s Horizon 2020 Research and Innovation Program under the Marie Skłodowska-Curie grant agreement 748627 and BMBF, RAPID Consortium, grant 01KI1723D. K.B.M. and J.W.S. were supported in part by The Clayton Foundation and The American Lung Association. J.W.S. was additionally supported by NIH grant AI117922 and a Burroughs Wellcome Fund ‘Investigators in the Pathogenesis of Infectious Diseases’ Award. NH was supported by NIH grant AI132751. E.M., H.H.H., M.S. and C.M.R. were supported in part by NIH grant AI091707. G.Z. was supported by a grant from the Swiss National Science Foundation grant 166265. VT was supported by the Swiss National Science Foundation (SNF; grants: 310030_173085; CRSII3_160780), and the Federal Ministry of Education and Research (BMBF; grant RAPID, #01KI1723A).

## Author contributions

S.P., K.B.M., E.M., C.M.R., J.W.S., and V.T. designed the project. S.P., K.B.M., E.M., A.K., D.H., P.V., W.F., N.E., H.S., H.K-W., M.H., H.H.H., M.S., M.W-C., N.W.H. and G.Z. performed experiments. R.D., S.Pö., E.S., and T.G. contributed to the design and implementation of the research. D.T. performed statistical analysis and analysis of RNA-Seq dataset. S.P., K.B.M., and J.W.S. wrote the manuscript, and all authors contributed to editing.

## Competing interest declaration

The authors declare no competing interests.

**Supplementary Information** is available for this paper.

## Materials & Correspondence

Correspondence and requests for materials should be addressed to Volker Thiel, John W. Schoggins, and Charles M. Rice.

## Extended Data

**Extended Data Figure 1.**
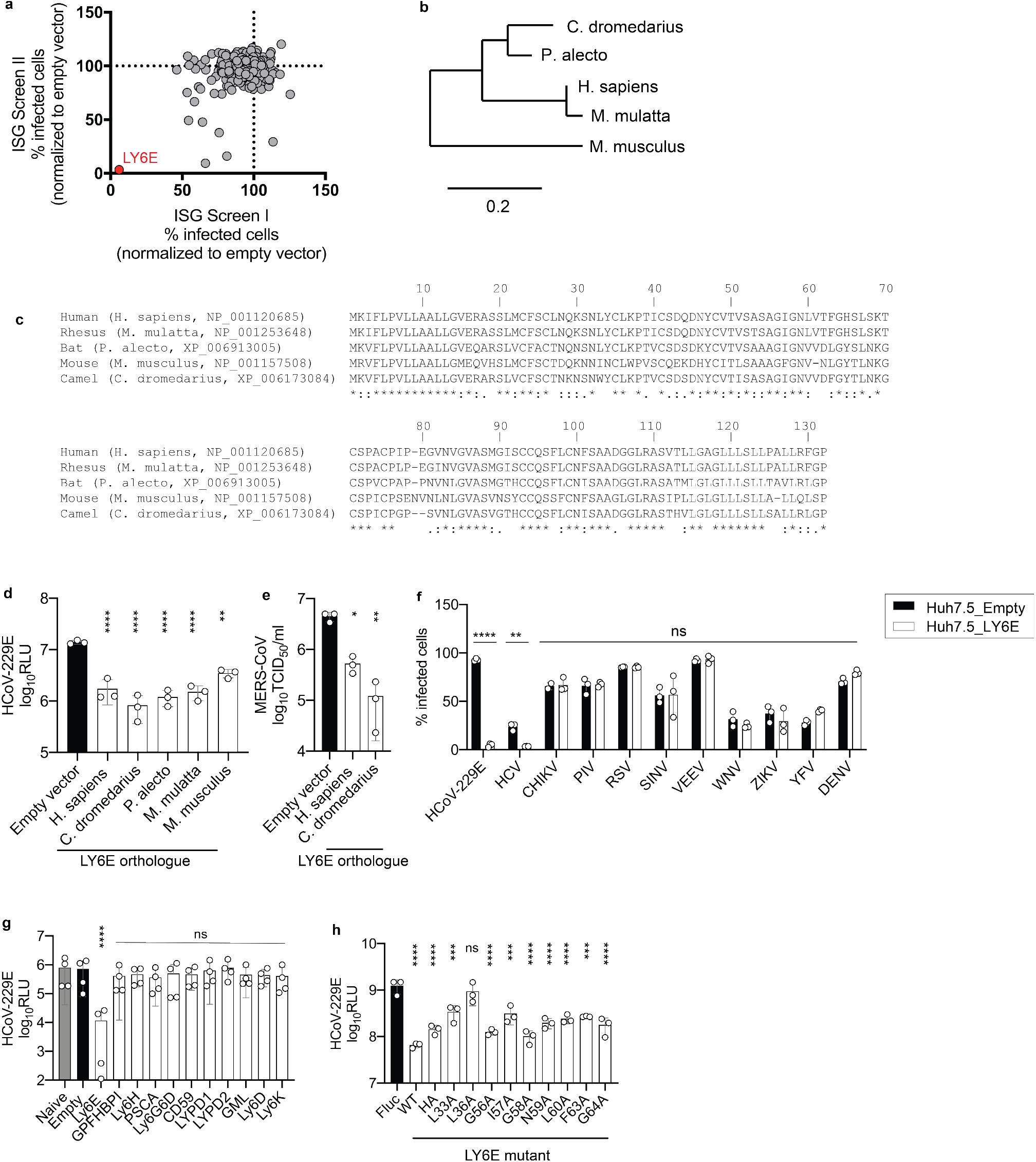
The antiviral activity of LY6E is evolutionarily conserved and specific to CoVs. **a**, Duplicate screens depicting HCoV-229E infection (48 hours post-infection) in Huh7 cells expressing ISGs. **b**, Multiple sequence alignment of Ly6e amino acid sequences of different species. **c**, Phylogenetic tree of mammalian LY6E orthologues. **d**, HCoV-229E infection of Huh7.5 cells expressing vector control or human (*H. sapiens*), camel (*C. dromedarius*), bat (*P. alecto*), mouse (*M. musculus*), or rhesus (*M. mulatta*) LY6E orthologues. **e**, Stable LY6E or empty vector expressing Huh7.5 cells infected with MERS-CoV in a limiting dilution assay. **f**, Stable Huh7.5 cells expressing either LY6E or containing empty vector infected with HCoV-229E, HCV, CHIKV, hPIV-3, RSV, SINV, VEEV, WNV, ZIKV, YFV, and DENV. **g**, HCoV-229E-Rluc infection of naïve or stably transduced Huh7 cells expressing empty vector control or LY6/uPAR family members. **h**, HCoV-229E-Rluc infection of Huh7 cells transduced with lentiviruses expressing firefly luciferase (Fluc) control, LY6E WT, LY6E HA or specific block mutants^9^. Data represent average of independent biological replicates, n=3 (**d**), n=3 (**e**), n=3 (**f**), n=3 (**g**), n=3 (**h**). Statistical significance was determined by ordinary one-way ANOVA with Dunnett’s correction for multiple comparison (**d, e, g, h**), 2way ANOVA followed by Sidak’s multiple comparisons test (**f**). Error bars: SD. *P* values: **d**, **** p=<0.0001, ** p=0.0030; **e**, * p=0.0118, ** p=0.0013; **f**, **** p=>0.0001, ** p=0.0095, ns= >0.9999 (CHIKV) >0.9999 (PIV) >0.9999 (RSV) >0.9999 (SINV) >0.9999 (VEEV) 0.9679 (WNV) 0.8794 (ZIKV) 0.3172 (YFV) 0.7667 (DENV) (For CHIKV one outlier was removed in the control cells); **g**, **** p=<0.0001, ns p=0.9993 (GPFHBPI) 0.9998 (LY6H) 0.9991 (PSCA) 0.9990 (Ly6G6D) 0.9997 (CD59) >0.9999 (LYPD1) 0.9994 (LYPD2) 0.9997 (GML) 0.9997 (LY6D) 0.9992 (LY6K); **h**, **** p=<0.0001, <0.0001, *** p=0.0005, ns p=0.8472, **** p=<0.0001, *** p=0.0003, **** p=<0.0001, <0.0001, <0.0001, *** p=0.0001, **** p=<0.0001.

**Extended Data Figure 2.**
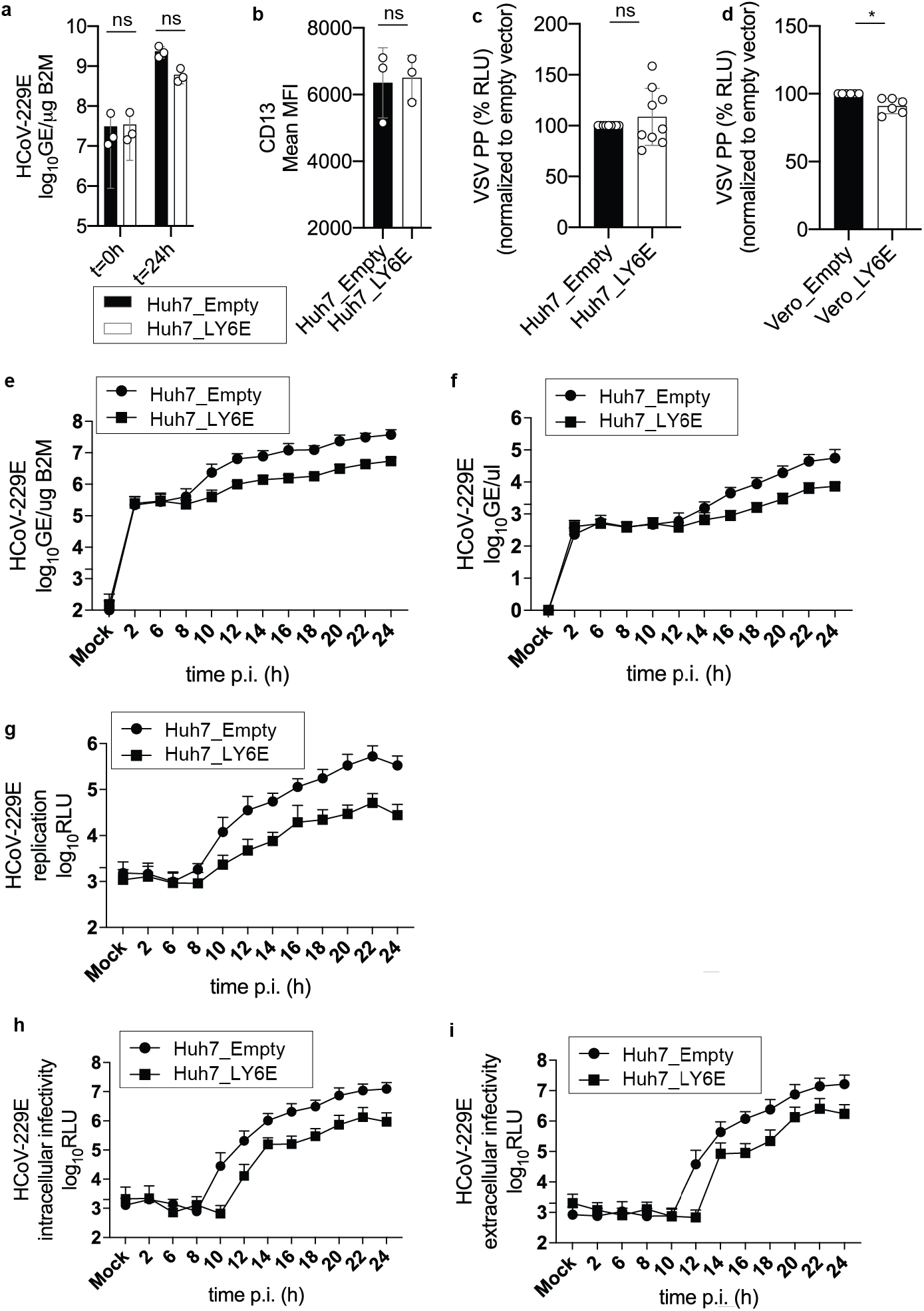
LY6E does not affect CoV binding, replication, translation, assembly, or release. **a**, Stable LY6E expressing or empty control Huh7 cells were incubated with HCoV-229E and binding analyzed immediately (t=0 h) or after incubation for 24 hours (t=24 h) by RT-qPCR. **b**, Stable LY6E or empty vector expressing Huh7 cells were assessed for surface CD13 expression by flow cytometry. **c**,**d,** VSV pseudoparticles (PP) harboring VSV-G were inoculated on stable LY6E or empty vector Huh7 cells (**c**) or VeroE6 cells (**d**). **e-i**, Stable LY6E expressing or control Huh7 cells were mock-infected or infected with HCoV-229E-Rluc. Cell lysates were harvested at the indicated time points and intracellular viral RNA was extracted, and viral replication detected via qRT-PCR (**e**). Cell supernatant was harvested and extracellular viral RNA was extracted and viral replication detected via qRT-PCR (**f**). Cell lysates were harvested and intracellular Renilla luciferase activity was detected upon cell lysis (**g**). Cells were subjected to 3 rounds of freeze/thaw cycles. Cell debris was removed and the supernatant titrated on naïve Huh7 cells. Intracellular infectivity was determined (**h**). Supernatant was harvested and titrated on naïve Huh7 cells to determine extracellular infectivity (**i**). Data represent averages of independent biological replicates, n=3 (**a**), n=3 (**b**), n=9 (**c**), n=6 (**d**), n=3 (**e-i**). Statistical significance was determined by 2way ANOVA followed by Sidak’s multiple comparisons test (**a**), unpaired student’s t-test with Welch’s correction (**b, c, d**). Error bars: SD. *P* values: **a**, ns p=0.9448 0.0752; **b**, ns p=0.5188; **c**, ns p=0.3857; **d**, * p=0.0112.

**Extended Data Figure 3.**
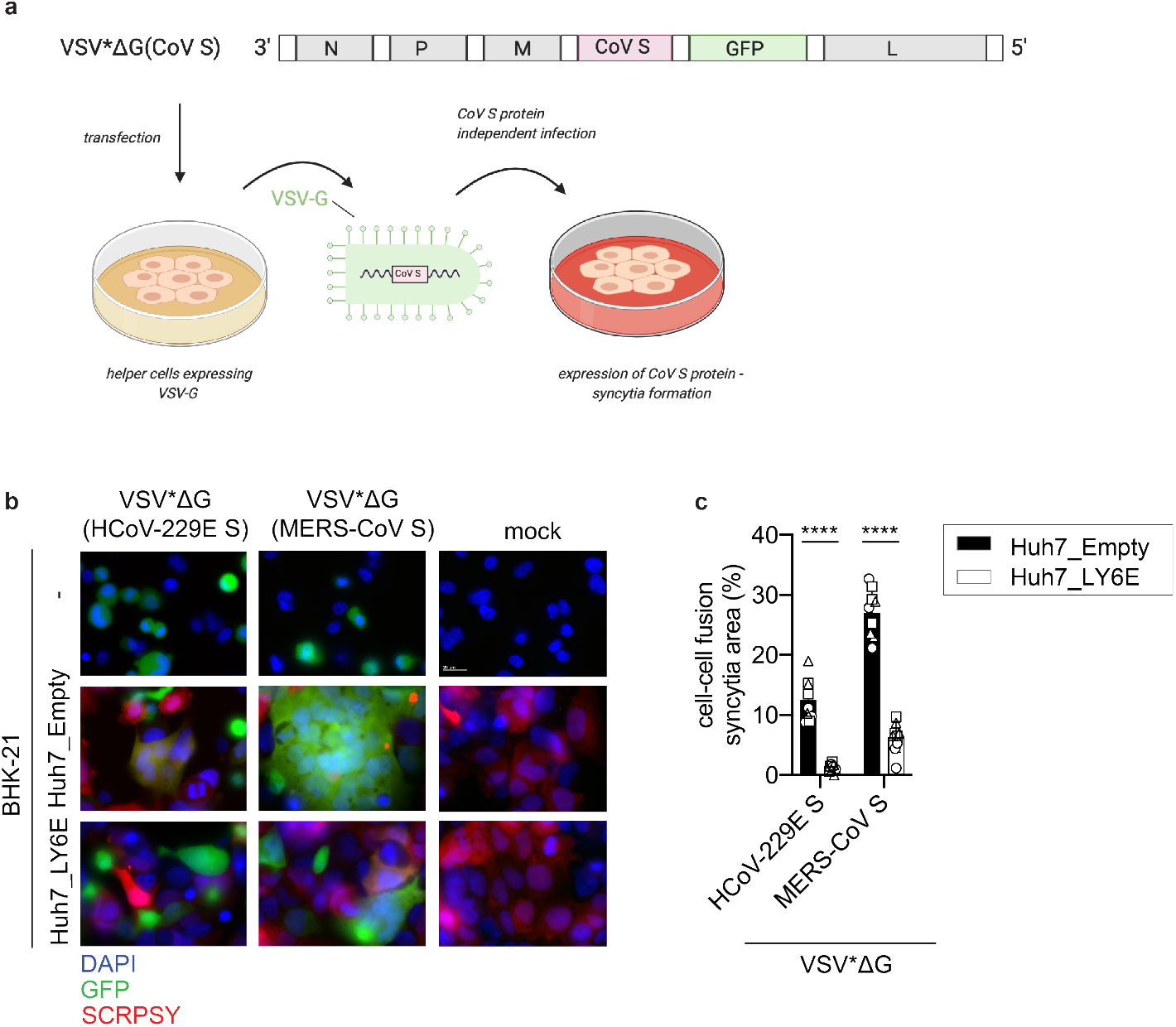
Generation of recombinant VSV vector driving CoV S protein expression (VSV*ΔG(CoV)) and heterologous cell-cell fusion assay. **a**, Schematic depiction of the generation of recombinant VSV*ΔG(CoV S) expressing both CoV S protein and GFP reporter protein. The respective CoV S genes were inserted into the genomic VSV*ΔG cDNA and recombinant virus was generated in BHK-G43 cells expressing the VSV G protein. The recombinant virus produced in this way harbored the VSV G protein in the envelope allowing CoV S protein-independent infection of cells. Infection of cells with VSV*ΔG(CoV S) led to the expression of CoV S protein and consequent syncytia formation. **b**, Heterologous syncytia formation assay. BHK-21 cells were infected with VSV G protein trans-complemented VSV*ΔG(CoV S) viruses or were mock-infected, followed by co-culture with LY6E or empty control Huh7 cells. Syncytia formation was determined. Blue: DAPI, green: GFP, red: TagRFP encoded in SCRPSY vector. **c**, Quantification of VSV*ΔG(CoV S) induced syncytia depicted as percentage syncytia area. Three independent areas were analyzed per biological replicate (circle, square, triangle). Data represent averages of independent biological replicates, n=3 (**c**). In **b**, scale bar is 20 µM. Statistical significance was determined by 2way ANOVA followed by Sidak’s multiple comparisons test (**c**). Error bars: SD. *P* values: **c**, **** p=<0.0001 <0.0001.

**Extended Data Figure 4.**
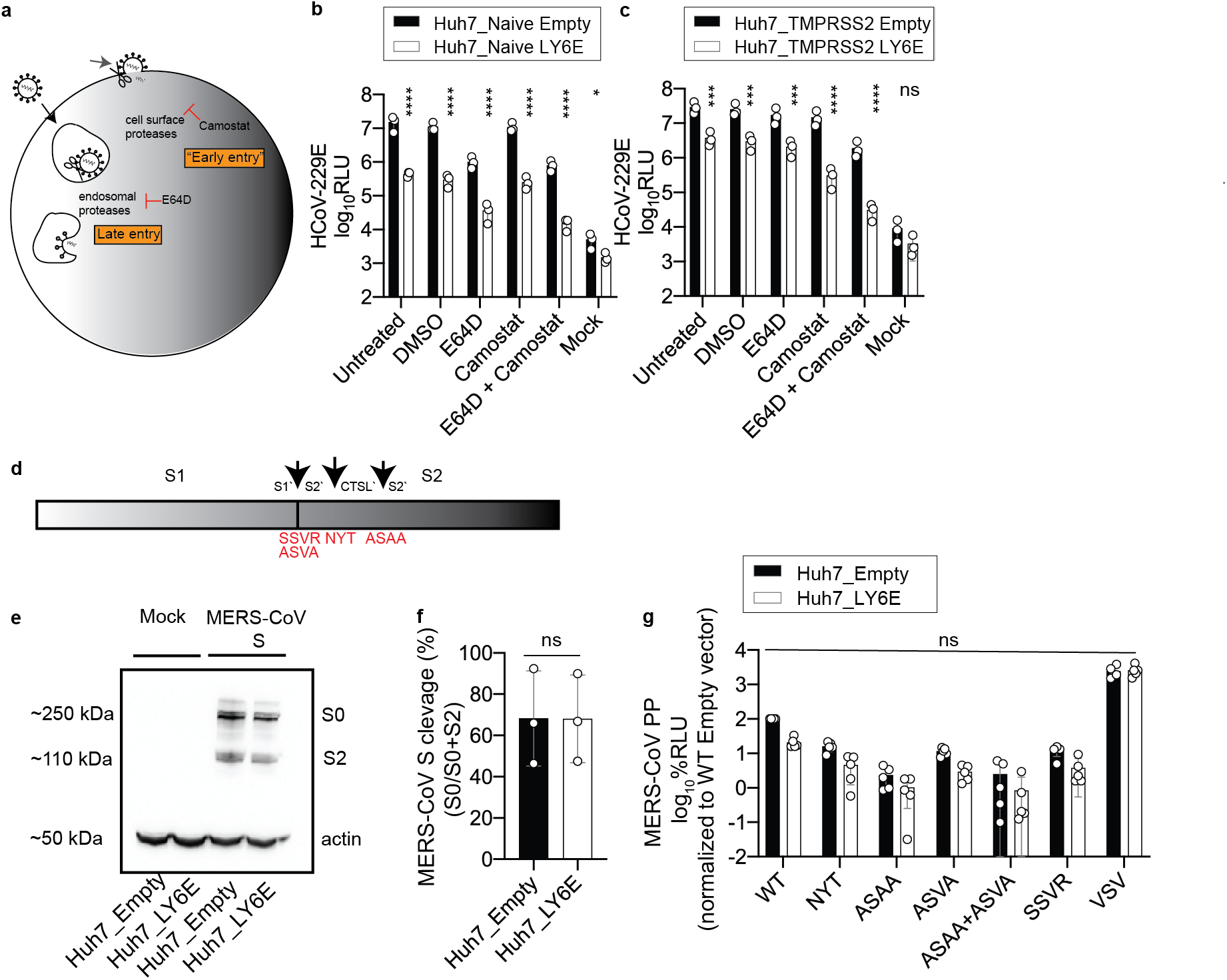
The antiviral effect of LY6E is independent of proteolytic cleavage of CoV spike protein. **a**, Schematic depiction of cell entry routes of CoVs and intervention by selected compounds. **b-c**, LY6E or empty control-expressing Huh7 cells naïve for (**b**) or ectopically overexpressing TMPRSS2 (**c**) were pre-treated with the indicated compounds before infection with HCoV-229E-Rluc. **d**, Schematic depiction of the CoV spike (S) protein containing the subunits S1 and S2. Arrows indicate the S1/S2’ cleavage site, a proposed cleavage site for cathepsin L (CTSL’) and the S2’ cleavage site. Amino acid exchanges disrupting the respective cleavage sites are depicted in red^17^. **e**, Western blot of S cleavage in LY6E or empty control-expressing Huh7 cells transfected with a plasmid encoding for MERS-CoV S protein (S0= uncleaved, S2= S2 subunit). **f**, MERS CoV S cleavage was analyzed by quantification of S0 and S2 bands. **g**, LY6E or empty control-expressing Huh7 cells inoculated with CoV-pseudoparticles (PP) harboring MERS-CoV S WT or MERS-CoV S proteins containing various cleavage site mutations. Data represent average of independent biological replicates, n=3 (**b**), n=3 (**c**), n=3 (**e**), n=3 (**f**), n=5 (**g**). Statistical significance was determined by 2way ANOVA followed by Sidak’s multiple comparisons (**b-c, g**), unpaired student’s t-test with Welch’s correction (**f**). Error bars: SD. *P* values: **b**, **** p=<0.0001, <0.0001, <0.0001, <0.0001, <0.0001, * p=0.0179; **c**, *** p=0.0007, 0.0005, 0.0006, **** p=<0.0001, <0.0001, ns p=0.1909; f, ns p=0.4968; **g**, ns p=0.9999 (WT) >0.9999 (NYT) >0.9999 (ASAA) >0.9999 (ASVA) >0.9999 (ASAA+ASVA) >0.9999 (SSVR) 0.9867 (VSV).

**Extended Data Figure 5.**
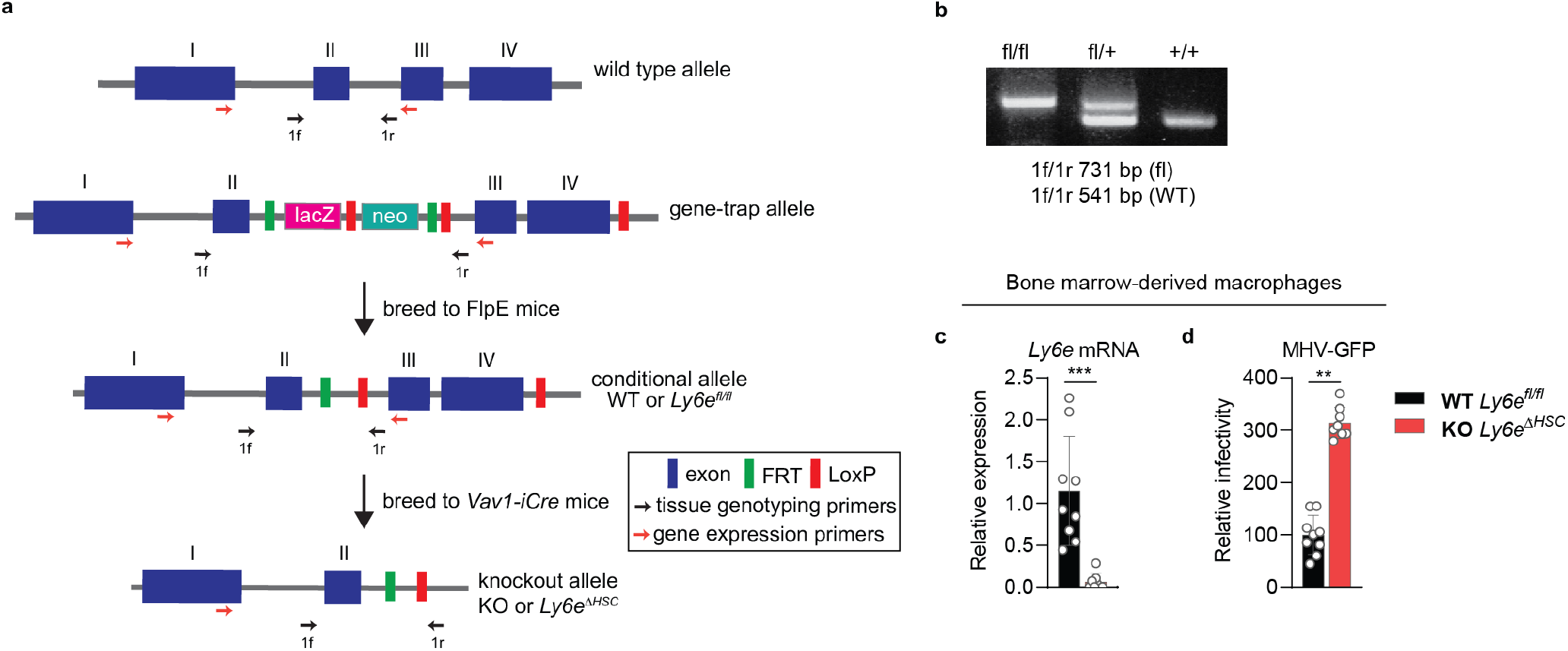
Generation of hematopoietic Ly6e conditional knockout mouse. **a,** Strategy used to generate *Ly6e^ΔHSC^* mice. Hematopoietic-tissue specific ablation of Ly6e was achieved by crossing *Ly6e^fl/fl^* mice with transgenic *Vav1-iCre* mice to remove the LoxP-flanked exon III and IV, resulting in a 17 amino acid truncated variant of the 130 amino acid full-length protein. Primers used for tissue genotyping and gene expression are also depicted as arrows. **b,** Gel electrophoresis of tissue genotyping PCR, representing *Ly6e^fl/fl^*, *Ly6e^fl/+^*, and *Ly6e^+/+^* mice. **c**, *Ly6e* gene expression in bone marrow-derived macrophages (BMDM). **d,** Infection of BMDM. For **c-d**, n=9 (*Ly6e^fl/fl^*) or n=8 (*Ly6e^ΔHSC^*) mice from three pooled experiments. Data for **c-d** is shown normalized to *Ly6e^fl/fl^* values. Statistical significance was determined by using two-tailed unpaired student’s t-test with Welch’s correction (**c**), two-tailed ratio-paired t-test (**d**). Error bars: SD. *P* values: **c**, *** p=0.0010; **d**, ** p=0.0021.

**Extended Data Figure 6.**
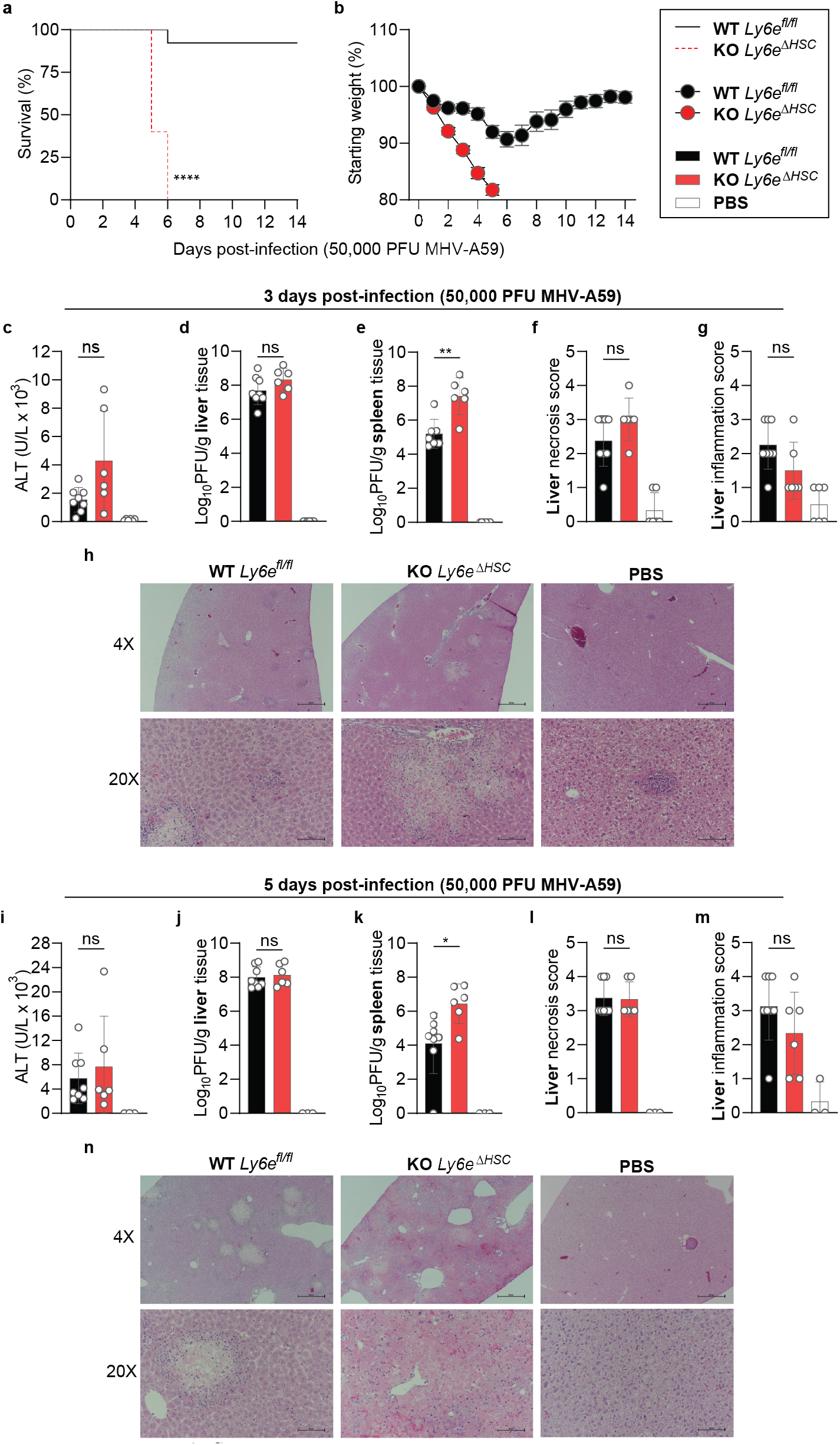
*Ly6e^ΔHSC^* mice are susceptible to murine hepatitis virus. **a-b**, Male *Ly6e^fl/fl^* and *Ly6e^ΔHSC^* mice injected with PBS or MHV and monitored for survival (**a**) and weight loss (**b**). **c-n,** Mice were euthanized at 3- (**c-h**) or 5- **(i-n**) days post-infection for determination of serum ALT (**c,i**), viral titers in liver (**d,j**) and spleen (**e,k**), and liver necrosis and inflammation (**f-h**, **l-n**). For **a-b**, n=15 for days 0-6, n=14 for days 6-14, (*Ly6e^fl/fl^*), n=10 for days 0-5, n=4 for days 5-6, n=0 for days 6-14 (*Ly6e^ΔHSC^*) from three pooled experiments. For **c-g**, n=8 (*Ly6e^fl/fl^*,), n=6 (*Ly6e^ΔHSC^*), or n=6 (PBS) from two pooled experiments. For **i-m**, n=8 (*Ly6e^fl/fl^*,), n=6 (*Ly6e^ΔHSC^*), or n=3 (PBS) from two pooled experiments. In **h,n**, scale bars are 500 µM (4x) and 100 µM (20x). Significance for was tested by Mantel-Cox test (**a**), two-tailed unpaired student’s t-test with Welch’s correction (**c-e**, **i-k**), two-tailed Mann-Whitney U test (**f-g**, **l-m**). Error bars: SEM (**b**), SD (**c-g**, **i-m**). *P* value: **a**, **** p=3.946 × 10^−6^; **c,** ns p=1144; **d**, ns p=0.1407; **e**, ** p=0.0026; **f**, ns p=0.1708; **g**, ns p=0.0779; **i**, ns p=0.6191; **j**, ns p=0.6842; **k**, * p=0.0119; **l**, ns p>0.9999; **m**, ns p=0.0779.

**Extended Data Figure 7.**
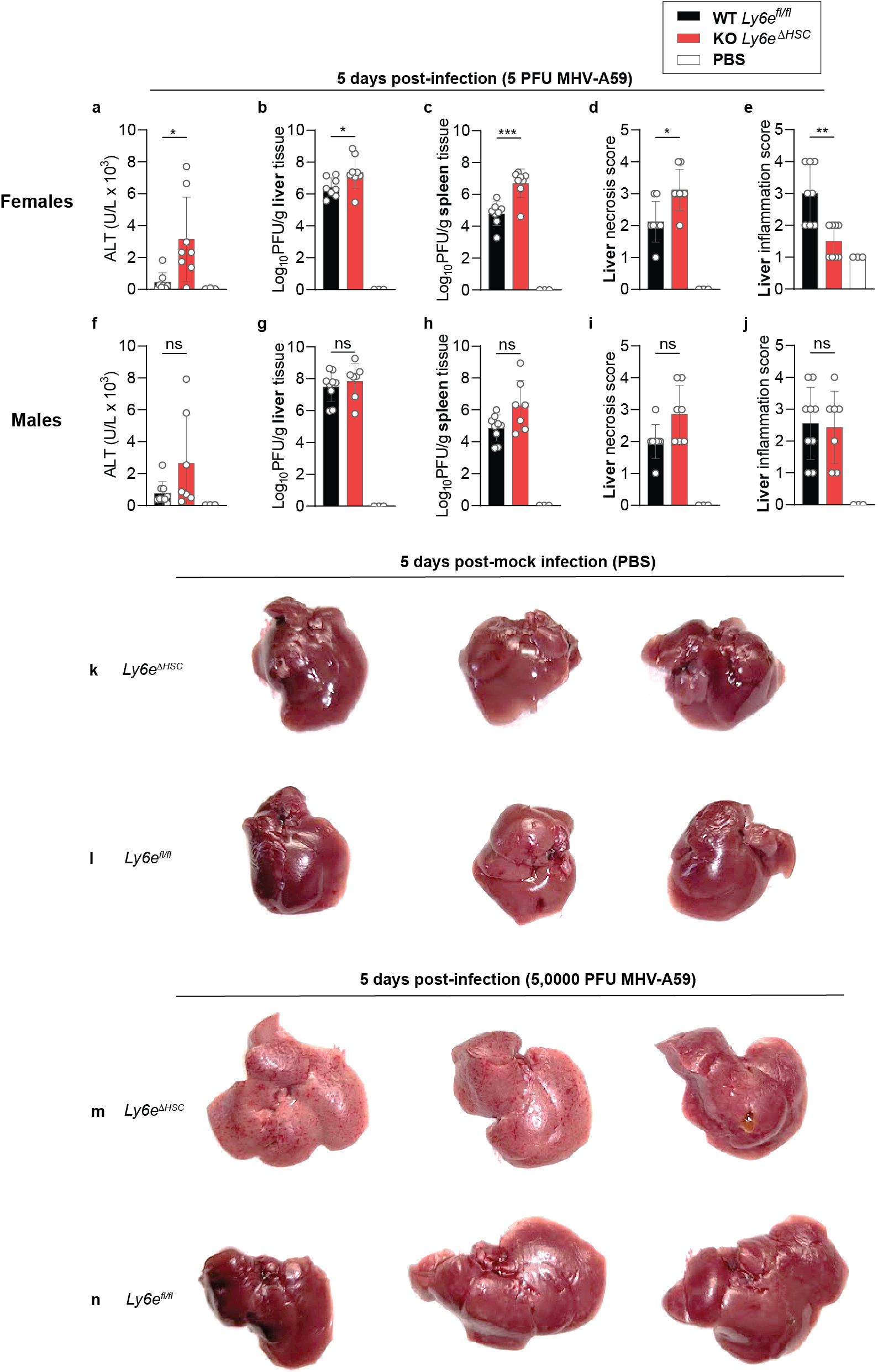
*Ly6e^ΔHSC^* mice have enhanced sensitivity to murine hepatitis virus. **a-j**, Female (**a-e**) and male (**f-g**) *Ly6e^fl/fl^* and *Ly6e^ΔHSC^* mice injected with PBS or MHV and assessed for serum ALT (**a**,**f**), viral titers in liver (**b**,**g**) and spleen (**c**,**h**), and liver necrosis and inflammation (**d-e**, **i-j**). **k-n**, Female *Ly6e^fl/fl^* and *Ly6e^ΔHSC^* mice injected with PBS or MHV and whole livers were imaged. Mock-infected (PBS) *Ly6e^ΔHSC^* mice (**k**), mock-infected *Ly6e^fl/fl^* mice (**l**), MHV-infected *Ly6e^ΔHSC^* mice (**m**), MHV-infected *Ly6e^fl/fl^* mice (**n**). For **a-e**, n=8 (*Ly6e^fl/fl^*,), n=8 (*Ly6e^ΔHSC^*), or n=3 (PBS) from two pooled experiments. For **f-j**, n=9 (*Ly6e^fl/fl^*,), n=7 (*Ly6e^ΔHSC^*), or n=3 (PBS) two pooled experiments. For **k-n**, n=3. Statistical significance was determined by two-tailed unpaired student’s t-test with Welch’s correction (**a-c**, **f-h**), two-tailed Mann-Whitney U test (**d-e**, **i-j**). Error bars: SD. *P* value: **a**, * p=0.0234; **b**, * p=0.0428; **c**, *** p=0.0004; **d,** * p=0.0210; **e**, ** p=0.0054; **f**, ns p=0.1544; **g**, ns p=0.5322; **h**, ns p=0.0662; **i**, ns p=0.0592; **j**, ns p=0.9064.

**Extended Data Figure 8.**
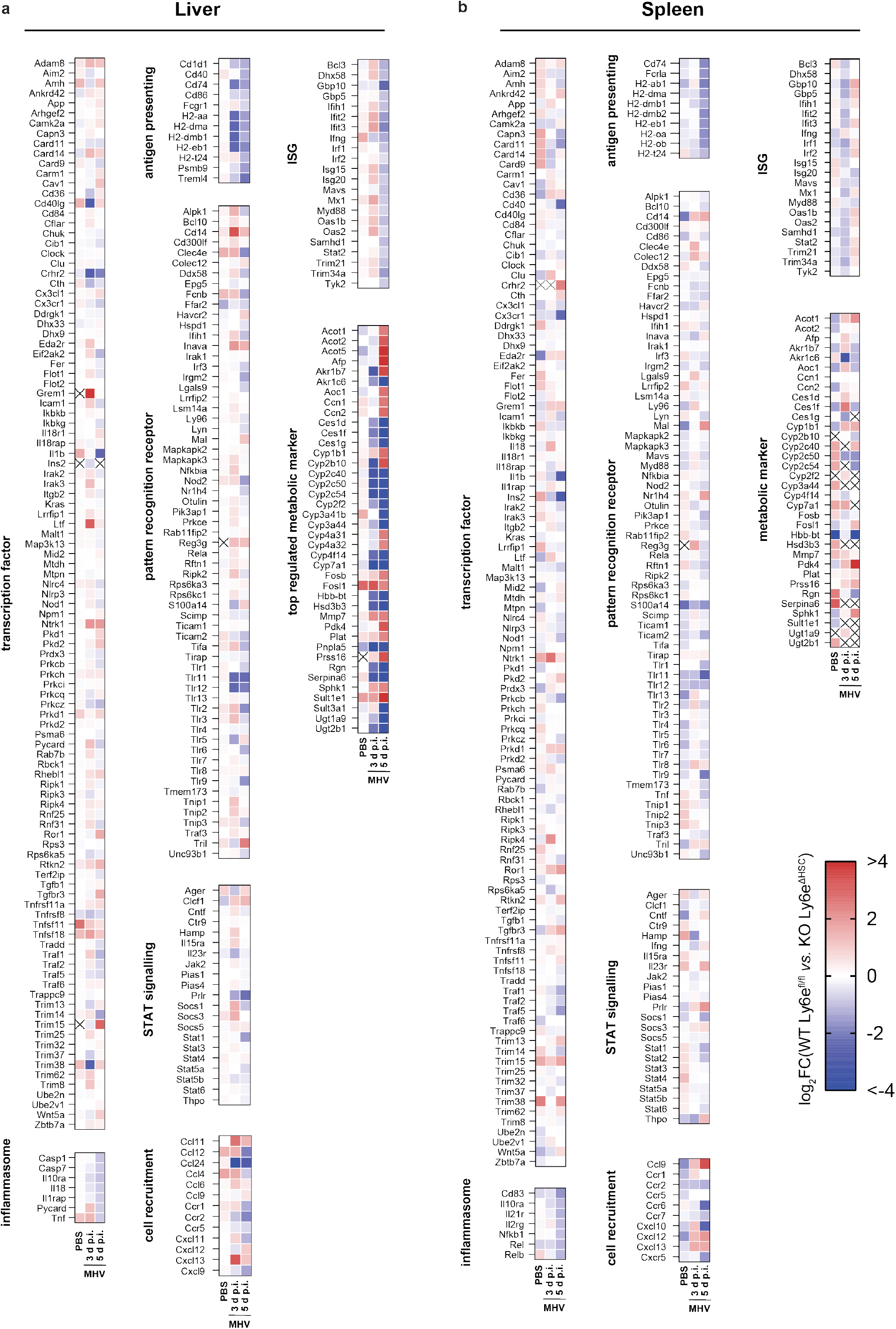
Dynamic transcriptional changes in liver and spleen of *Ly6e^ΔHSC^ vs Ly6e^fl/fl^* mice. Depicted are fold changes in liver and spleen gene expression from *Ly6e^ΔHSC^* mice compared to *Ly6e^fl/fl^* mice. RPKM values from RNAseq data for liver (**a**) and spleen (**b**) were used to calculate fold changes. Crossed out genes were not detected in either KO or WT mice at respective conditions.

**Extended Data Figure 9.**
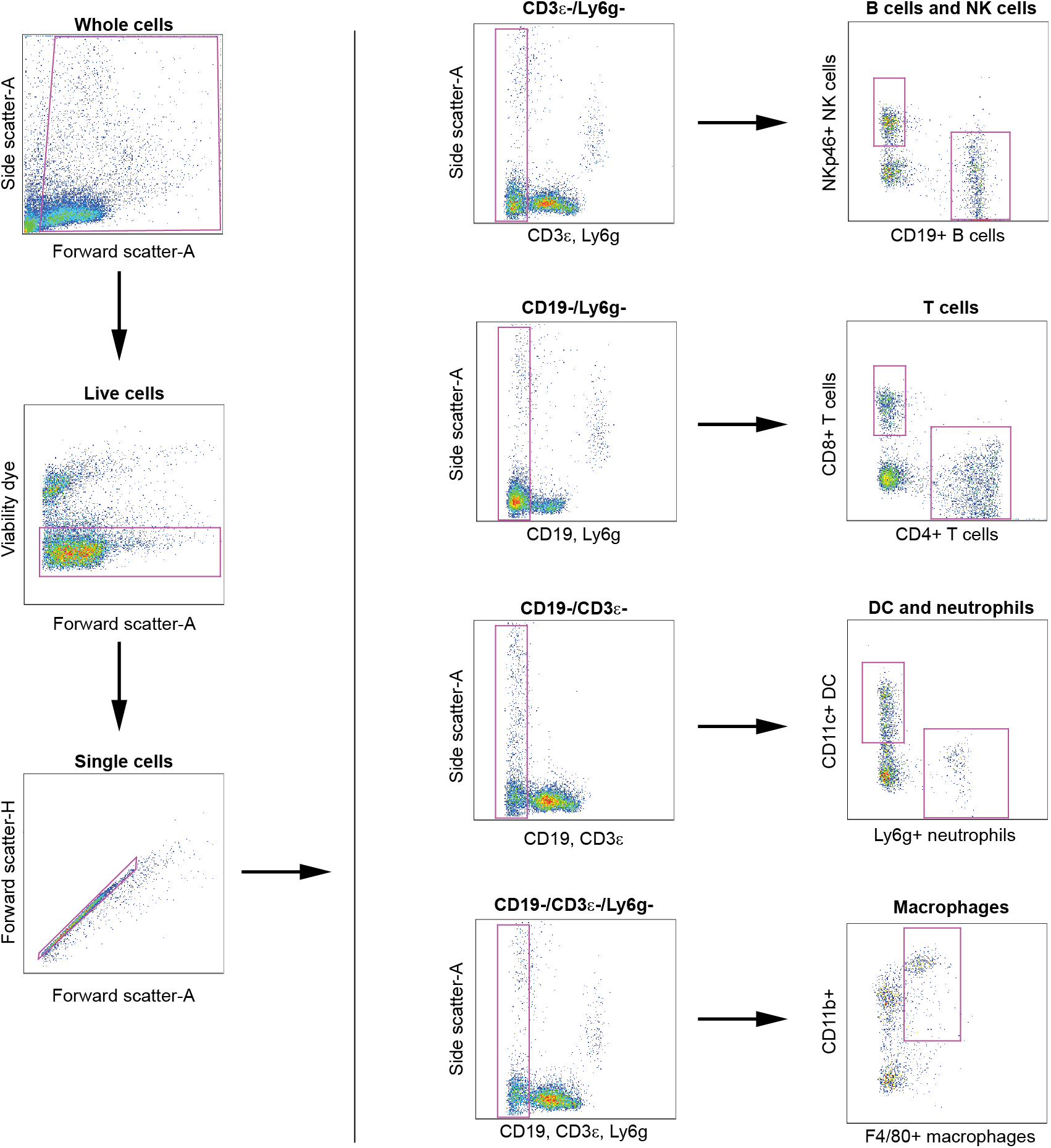
Gating strategy for intrahepatic and splenic immune cell identification. Antibody-stained suspensions of isolated intrahepatic immune cells or splenic immune cells were first analyzed by forward scatter/side scatter to remove debris events, then gated for exclusion of a viability dye, and finally for selection of single cell events. A dump-gate strategy was used to identify immune cell populations as indicated in the diagram.

**Extended Data Figure 10.**
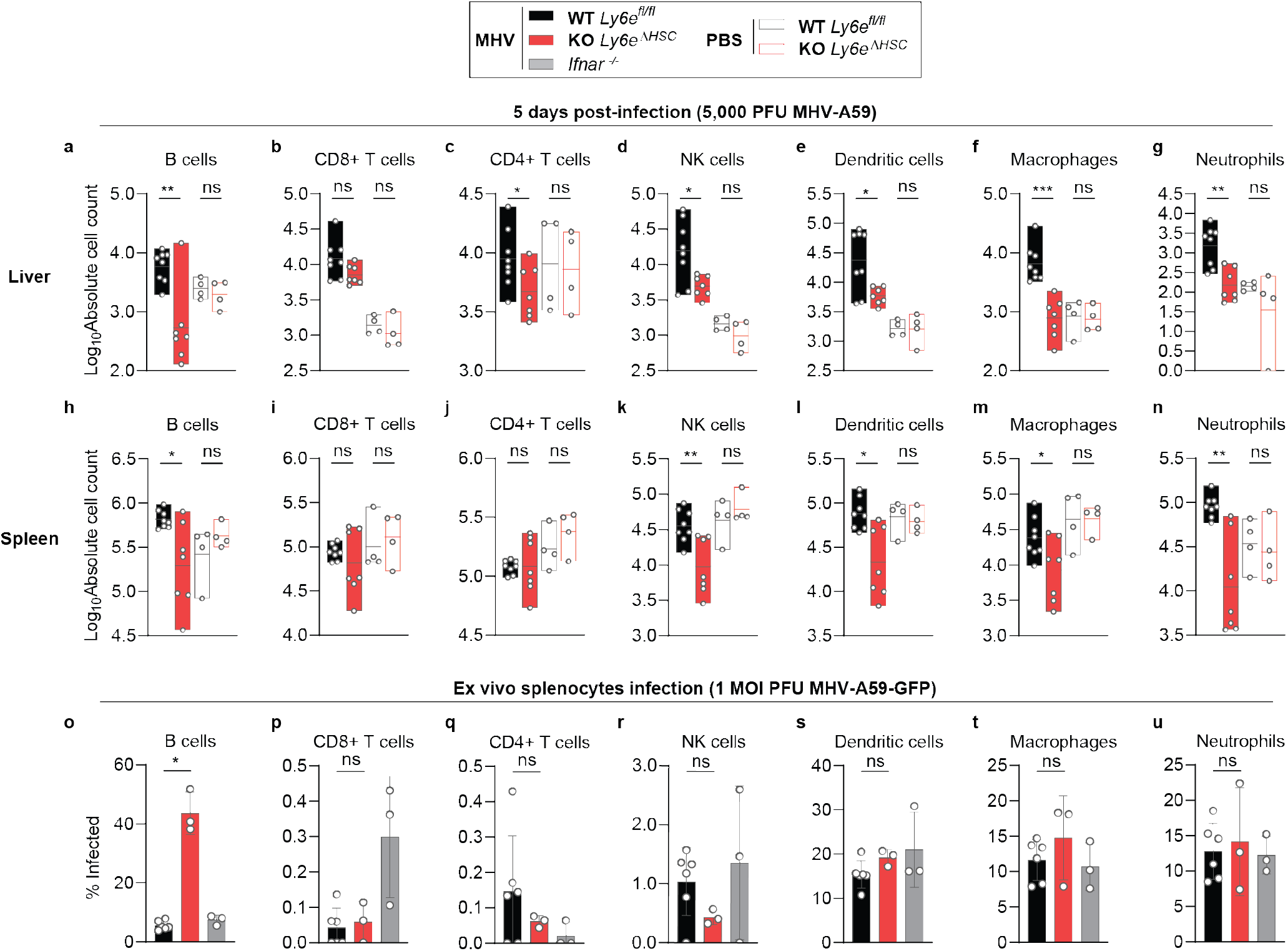
Murine hepatitis virus infection significantly alters hepatic and splenic immune cell populations in male *Ly6e^ΔHSC^* mice. **a-b**, Immune cell counts from liver (**a**) and spleen (**b**). **c**, Infection of cultured splenocytes. For **a-b**, n=8 (infected *Ly6e^fl/fl^*), n=7 (infected *Ly6e^ΔHSC^*), n=4 PBS-injected from two pooled experiments. For **h**, n=6 (*Ly6e^fl/fl^*), n=3 (*Ly6e^ΔHSC^*), or n=e (*Ifnar^−/−^*) mice from two pooled experiments. Statistical significance was determined by two-tailed unpaired student’s t-test with Welch’s correction (**a-c**). Error bars: SD. *P* values (left-right): **a**, ** p=0.0053, ns p=0.5297, ns p=0.0641, ns p=0.4066, * p=0.0429, ns p=0.8678, * p=0.0163, ns p=0.2261, * p=0.0148, ns p=0.9694, *** p=0.0001, ns p=0.7840, ** p=0.0011, ns p=0.3465; **b**, * p=0.0284, ns p=0.3291, ns p=0.4421, ns p=0.6188, ns p=0.9412, ns p=0.2802, *** p=0.0082, ns p=0.4336, * p=0.0105, ns p=0.6763, * p=0.0419, ns p=0.9712, ** p=0.0046, ns p=0.6857; **c**, * p=0.0105, ns p=0.7080, ns p=0.2470, ns p=0.0503, ns p=0.0568, ns p=0.4637, ns p=0.7915.

**Extended Data Table 1.**
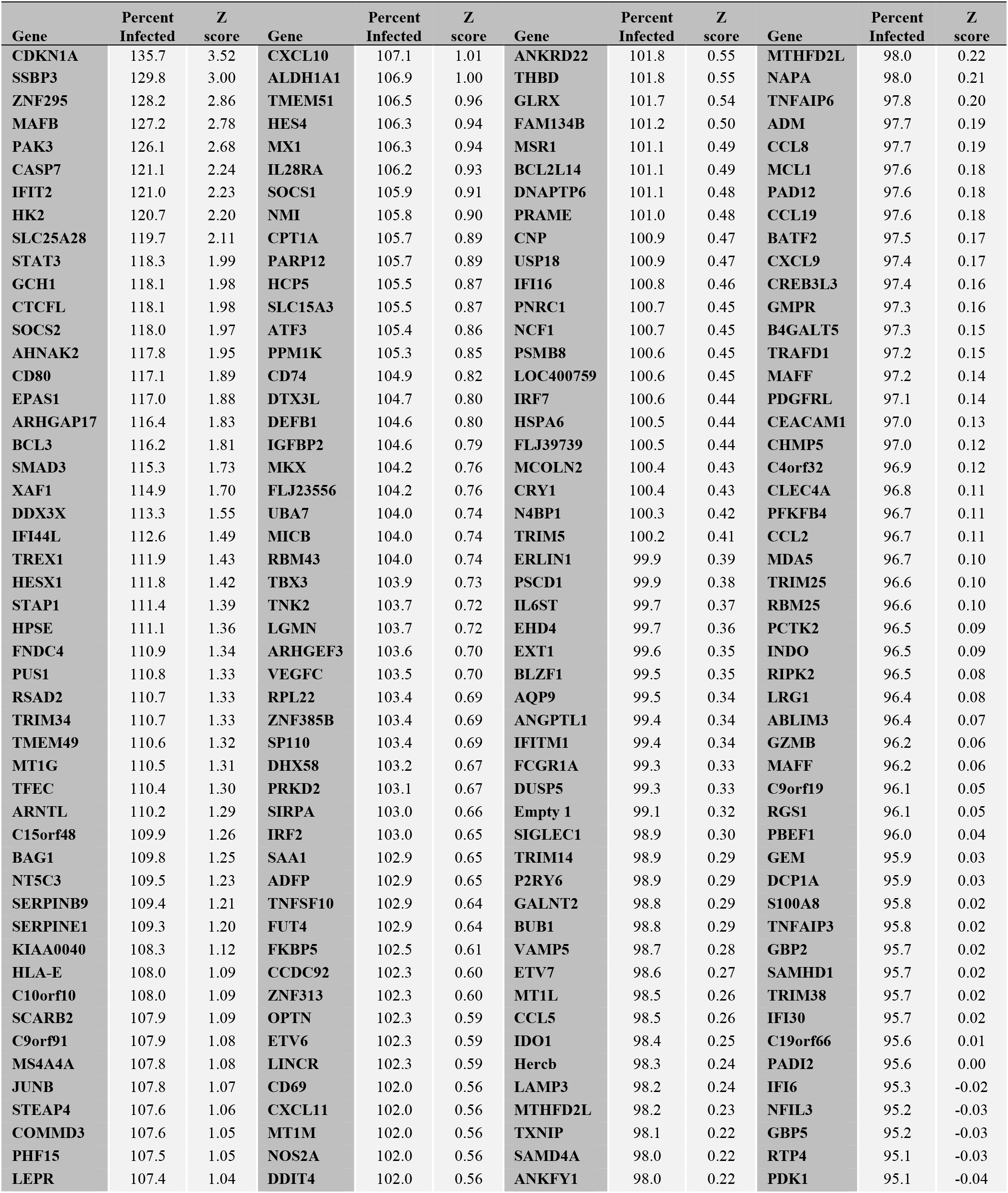
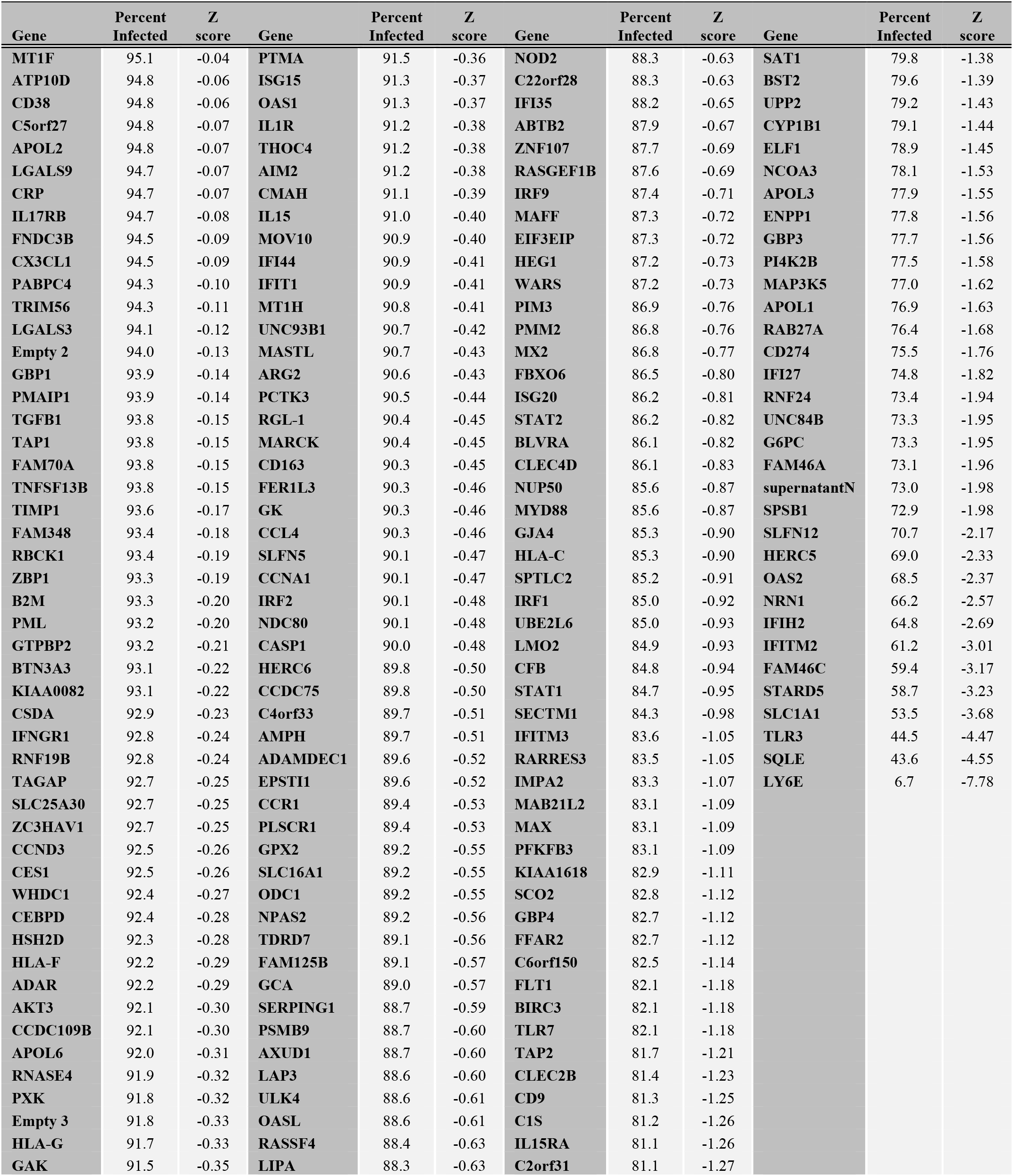
Large scale ISG screen: HCoV-229E in Huh7 at 24 h.

**Extended Data Table 2.**
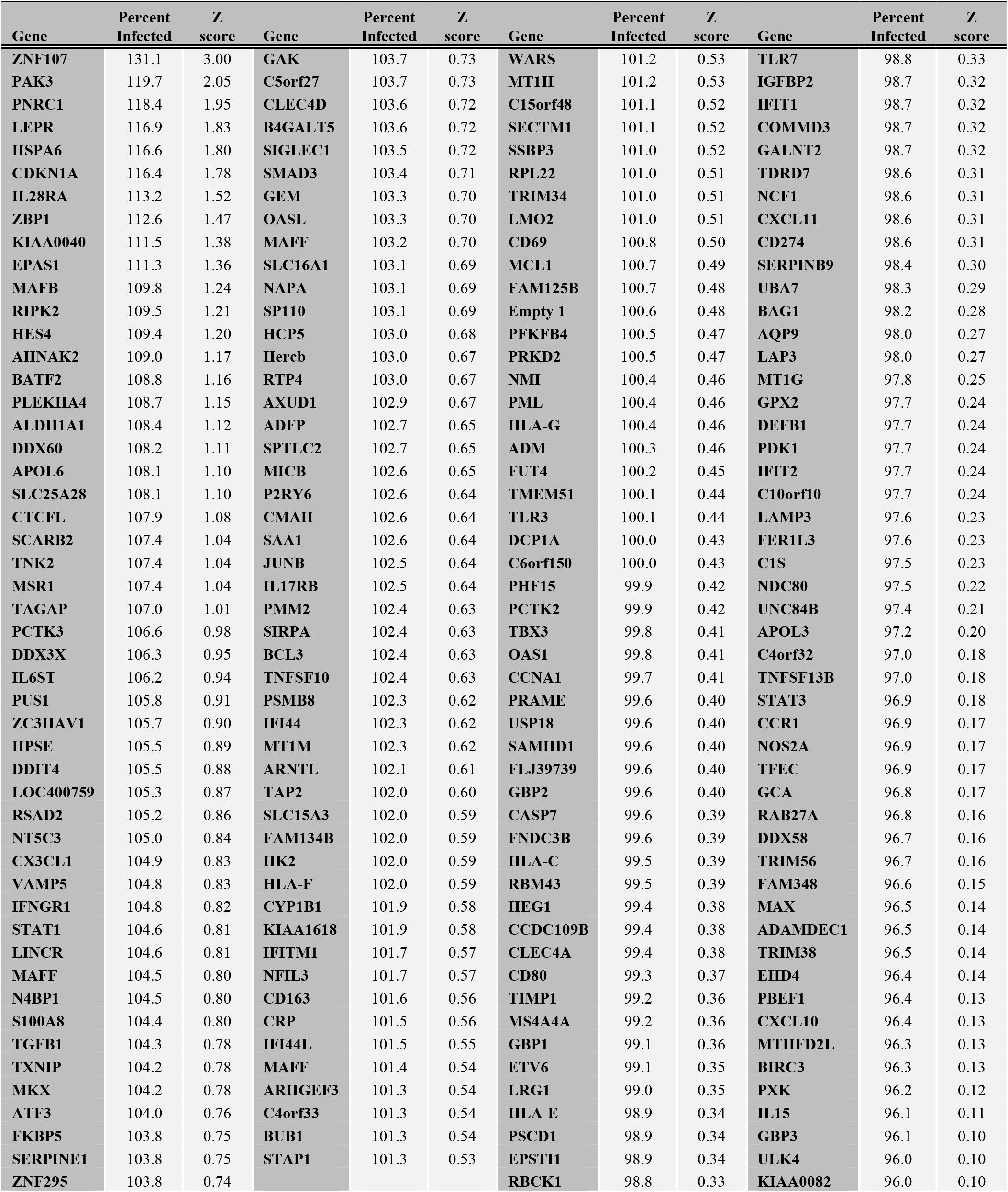
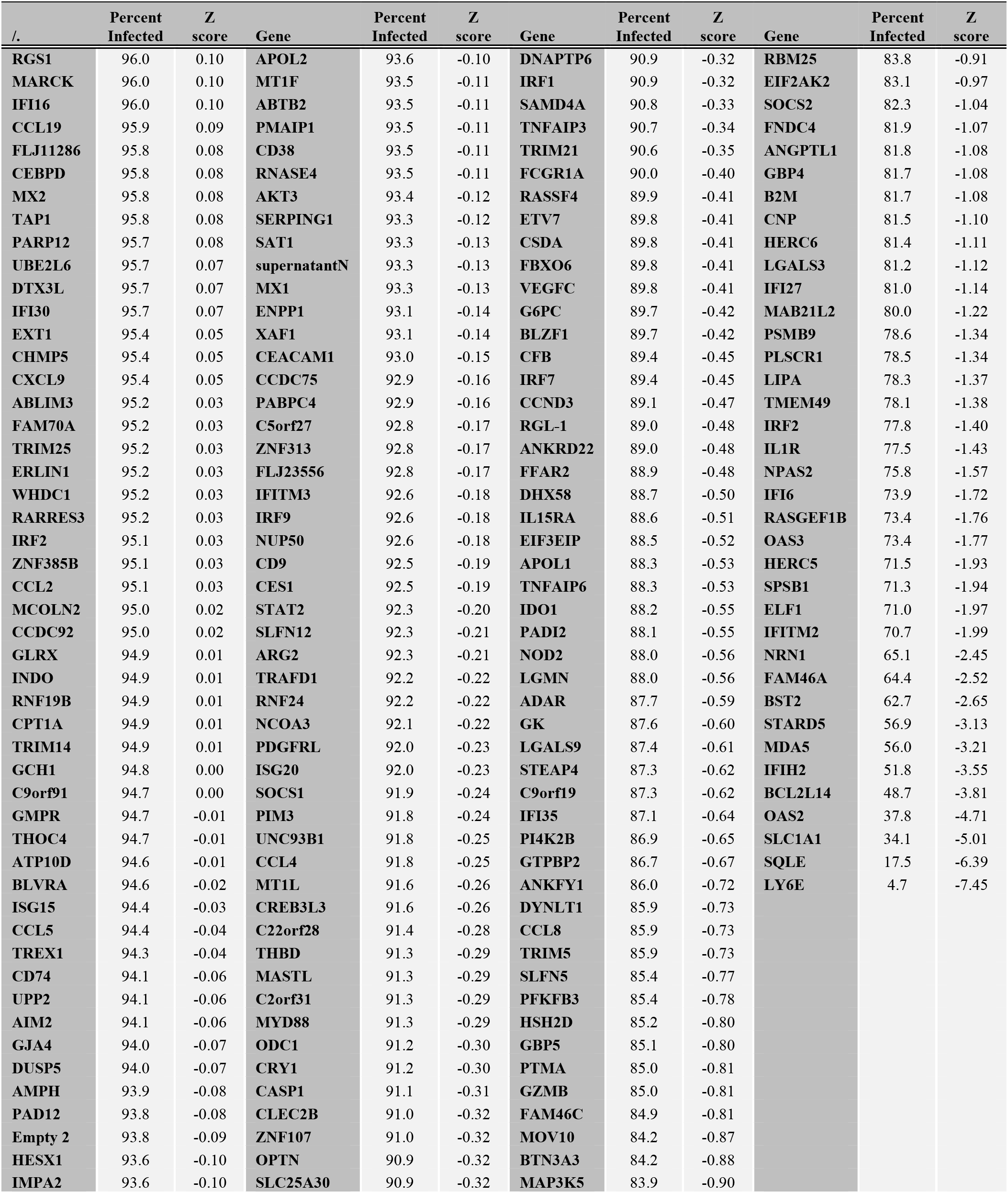
Large scale ISG screen: HCoV-229E in Huh7 at 48 h.

### Common ISG Aliases

DDX3X - DDX3, VIPERIN - RSAD2, SERPINE1 - PAI 1, IL28RA - CRF2, VEGFC – VRP, SP110 - IFI41, IRF2 - PRIC285, ZNF313 - RNF114, DNAPTP6 - LOC26010, LOC400759 – PSEUDOGENE, PSCD1 - CYTH1, IFITM1 - IFI17, SIGLEC1 – FRAG, CCL5 - SYCA5, CXCL9 – MIG, IFIH1 - MDA5, INDO – IDO, C9orf19 - GLIPR2, PBEF1 – NAMPT, FLJ11286 - C19orf66, TAP1 - ABCB2, IFIT1 - IFI56/ISG56, FER1L3 – MYOF, CCR1 - CMKBR1, TDRD7 - PCTAIRE2BP, SERPING1 - C1NH, ENPP1 - PDNP1, BST2 - THN

## Supplementary Information

### Methods

#### Cell lines

Huh7 hepatocarcinoma cells (kind gift from V. Lohnmann), Huh7.5, STAT1^−/−^ fibroblasts (kind gift from J.-L. Casanova), VeroE6 cells and VeroB4 cells (kindly provided by M.Müller/C.Drosten), 293LTV (Cell Biolabs), and A549 cells (ATCC cat# CCL-185) were maintained in Dulbecco’s Modified Eagle Medium-GlutaMAX (Gibco) supplemented with, 1 mM sodium pyruvate (Gibco), 10% (v/v) heat-inactivated fetal bovine serum (FBS) (Gibco), 100 µg/ml streptomycin (Gibco), 100 IU/ml penicillin (Gibco) and 1% (w/v) non-essential amino acids (NEAA; Gibco) (cDMEM) BHK-21 (ATCC cat# CCL-10) were maintained in Glasgow’s minimal essential medium (MEM) with 5% FBS and 1% tryptose. BHK-G43^57^ were cultured in Glasgow’s minimal essential medium with 5% FBS. 17Cl1 fibroblasts (gift from S.G. Sawicki) were cultured in MEM supplemented with 10% (v/v) heat inactivated FBS, 100 µg/ml streptomycin and 100 IU/ml penicillin. Cells were either newly purchased or cell line identities (human cells only) verified by a Multiplex human cell line authentication test. Huh7 cells expressing TMPRSS2 were generated as follows. In order to generate TMPRSS2-encoding retroviral vectors, the open reading frame of human TMPRRS2 was first PCR amplified with primers adding an N-terminal cMYC epitope to the TMPRSS2 coding sequence. The resulting sequence was inserted into a modified version of the pQCXIP plasmid that contains a blasticidin resistance cassette instead of the usual puromycin resistance cassette^17^. Huh7 cells stably expressing human TMPRSS2 were generated by retroviral transduction and selection with the antibiotic blasticidin (50 µg/ml). Following selection, cells were maintained in culture medium (cDMEM) supplemented with 10 µg/ml blasticidin. To generate STAT1^−/−^_CEACAM1 cells, human STAT1^−/−^ fibroblasts were transduced with lentiviruses encoding for the murine CoV receptor CEACAM1 (kind gift from David Wentworth, CDC, Atlanta, USA^59^ and subsequently selected with 1 µg/mL puromycin. Stable LY6E expressing cell lines were generated upon lentiviral transduction with SCRPSY LY6E or as control SCRPSY Empty in DMEM containing 4 µg/ml polybrene (Millipore) and 20 mM HEPES buffer solution (Gibco). Cells were selected using 2-4 µg/mL puromycin and were passaged until all cells in control wells without lentivirus were killed. The puromycin selected cells were further passaged and frozen down for subsequent experiments. The stable cell lines harboring LY6E orthologues have been described before, with the exception of *C. dromedarius*^9^. For this, a gBlock was ordered (XM_031439745.1) and Gateway cloning performed as described previously to generate pENTR and pSCRPSY plasmids^9^. Constructs encoding for Ly6/uPAR family members and LY6E ASM mutants have been described before^9^.

To generate clonal LY6E KO cells and CRISPR-resistant LY6E, A549 cells were transduced with lentivirus containing a LY6E-specific sgRNA and Cas9 as described previously^9^. To generate a clonal cell line, the bulk transduced population was plated at single cell dilutions. Candidate clones were screened by Western blot for LY6E expression and Sanger sequencing. Silent mutations in the region targeted by the LY6E-specific sgRNA were introduced into HA-tagged LY6E to generate CRISPR-resistant LY6E (CR LY6E). LY6E KO A549 cells were reconstituted with CR-LY6E by lentiviral transduction and expression confirmed by Western blot.

All cell lines were regularly tested to check they were free of mycoplasma contamination using a commercially available system (PCR Mycoplasma test kit I/RT Variant C, PromKine).

#### ISG screen

The ISG screen was performed as described previously with slight modifications^7,8^. Briefly, 5 × 10^3^ Huh7 cells were seeded, transduced with individual lentiviruses, and 48 hours post-transduction infected with HCoV-229E at 33°C. Infection was stopped 24 hours (MOI=0.1) or 48 hours (MOI=0.01) post-infection and plates were immuno-stained as described previously^61^ with an anti-HCoV-229E N protein antibody and a AlexaFluor488-conjugated donkey anti-mouse secondary antibody (Extended methods; Antibodies for immunofluorescence and flow cytometry). For high-content high-throughput imaging analysis ImageXpress Micro XLS (Molecular Devices, Sunnyvale, CA) was used as previously described^8^. Hits were normalized to cells expressing the empty vector. Depicted are genes that were expressed in two independent screens with a transduction efficiency of at least 1 % (cut off). ISGs which were cytotoxic for the cells were excluded from the analysis. Normalized data can be found in **Extended Data Table 1** and **Extended Data Table 2**.

#### Western blotting

For SDS-PAGE and Western blot analysis, lysates from cultured cells were prepared using the M-PER Mammalian Protein Extraction Reagent (Thermo) supplemented with protease inhibitors (cOmplete Mini, Roche). Proteins were separated on 10% (w/v) SDS-polyacrylamide gels (Bio-Rad), and electroblotted on nitrocellulose membranes (in a Mini Trans-Blot cell (eBlot L1 GenScript). Membranes were incubated in PBS 0.1% Tween (Merck Millipore) 5% milk (Dietisa, Biosuisse), probed with the respective primary antibodies, followed by incubation with horseradish peroxidase-conjugated secondary antibodies. LY6E was detected using anti-LY6E rabbit monoclonal antibody GEN-93-8-1 (Genentech) at a dilution of 1:5,000 and secondary Peroxidase-AffiniPure Donkey Anti-Rabbit IgG (Jackson Immunoresearch, 711-035-152) at a dilution of 1:10,000. MERS-CoV spike was detected using the monoclonal anti-human 1.6c7 ab (0.12 mg/ml)^62^ (kindly provided by B.J. Bosch) at a dilution of 1:1,000 and a secondary Peroxidase-AffiniPure Donkey Anti-Human IgG (Jackson Immunoresearch, 709-035-098) at a dilution of 1:10,000. β-actin was detected using Monoclonal Anti-β-Actin-Peroxidase clone AC-15 (Sigma-Aldrich, A3854) at a dilution of 1:25,000. Proteins were visualized using WesternBright enhanced chemiluminescence horseradish peroxidase substrate (Advansta) and quantified in a Fusion FX7 Spectra (Vilber-Lourmat). Bands were quantified using the respective software (FusionCapt Software Version 18.05).

#### RNA extraction and RT-qPCR

Viral RNA was extracted from cell lysates (NucleoMag-96RNA kit, Macherey Nagel) or supernatant (NucleoMag-Vet kit, Macherey Nagel) using the KingFisher Flex robot according to the manufacturer’s recommendations. The commercially available TaqManTM Fast Virus 1-Step Master Mix (Applied Biosystems) was used for RT-qPCR with 2 µl of RNA input added to 8 µl of prepared mastermix per sample. Viral RNA was detected using HCoV-229E specific primers normalized to a qRT-PCR standard for HCoV-229E (contains the M gene of HCoV-229E^63^). Intracellular viral RNA was normalized to total RNA (determined via the housekeeping gene Beta-2-Microglobulin (B2M)). Cellular RNA was extracted from cell lysates of Huh7 cells using the Nucleospin RNA kit (Machery Nagel) according to the manufacturer’s recommendations, quantified via NanoDrop, and used to generate a standard curve. PCR conditions are available upon request.

#### Cellular CD13 expression

LY6E expressing or control cells were dissociated (TryPLE Express, Gibco) and enumerated. 3 × 10^5^ cells were stained in duplicate using Zombie Aqua™ Fixable Viability Kit (Biolegend) according to the manufacturer’s recommendations. Cells were resuspended in 0.1% FBS/PBS and stained for CD13 expression (See ‘Antibodies for immunofluorescence and flow cytometry’). Cells were washed with 0.1% FBS/PBS and resuspend in equal volumes. Cell suspensions were analyzed by flow cytometry using a FACS Canto (BD) and analyzed with FlowJo Software (Treestar).

#### Binding experiment

Stable LY6E expressing or control Huh7 cells were seeded in a 24-well plate (4 × 10^4^ cells) and mock infected or inoculated with HCoV-229E (MOI=5) in OptiMEM (Gibco) on ice for 1 hour. Cells were washed at least 3x with PBS and harvested immediately (t=0 h) or incubated at 33°C for 24 hours (t=24 h), before cell lysis and extraction of viral RNA as described above. Viral RNA was detected via RT-qPCR normalized to the housekeeping gene B2M as described above.

#### Time course experiment

8 × 10^4^ cells stable LY6E expressing or control Huh7 cells were seeded in 12-well plates and mock infected or infected with 229E-CoV-Rluc (MOI=0.1) for 2 hours at 33°C in OptiMEM. Cells were washed 3x with PBS and samples harvested at the indicated time points. To determine intracellular replication, cell lysates were collected and viral RNA extracted (NucleoMag-96RNA kit, Macherey Nagel), followed by RT-qPCR as described before, or cells were lysed using the Renilla Luciferase Assay System kit (Promega) according to the manufacturer’s recommendations and Rluc activity determined. To determine extracellular viral replication, viral supernatant was harvested, and viral RNA extracted (NucleoMag-Vet kit, Macherey Nagel), followed by RT-qPCR as described before. To determine intracellular infectivity, cells were subjected to 3 rounds of freeze/thaw cycles, centrifuged to remove debris (4,000 × *g* for 10 min), and the supernatant titrated on naïve Huh7 cells. To determine extracellular infectivity, supernatant was harvested at the indicated time points and titrated on naïve Huh7 cells. For titration, 1 × 10^4^ Huh7 cells were seeded in a 96-well plate and infected with 22 µl of virus containing supernatant and incubated at 33°C. Cells were lysed 24 hours post-infection and infectivity determined using the Renilla Luciferase Assay System kit (Promega) according to the manufacturer’s recommendations.

#### Protease inhibitor treatment

To test various protease inhibitors, 2 × 10^4^ naïve or TMPRSS2-expressing, control or LY6E cells were seeded in a 96-well plate. One day post seeding, cells were pre-treated with the following compounds for 1 hour in OptiMEM, at 37°C: DMSO (1:500, Sigma-Aldrich), E64 D (10 µM, Sigma-Aldrich), and/or Camostat (100 µM, Sigma-Aldrich). Cells were mock infected or infected with HCoV-229E-Rluc (MOI 0.1) in the presence of the inhibitors for 2 hours at 33°C. Medium was changed to DMEM and cells incubated for 24 hours at 33°C. Cells were lysed and Rluc activity detected using the Renilla Luciferase Assay System kit (Promega) according to the manufacturer’s recommendations.

#### Spike cleavage assays

LY6E expressing or control Huh7 (2 × 10^5^ cells) were seeded in 6-well cell culture plates. Cells were transfected using Lipofectamine 2000 (Invitrogen) with an expression plasmid encoding for MERS-CoV S (pCAGGS-MERS S). Cell lysates were harvested 48 hours post-transfection and subjected to Western blot analysis as described.

#### Multiple Sequence alignment and phylogeny

Multiple sequence alignment of LY6E amino acid sequences of different species (NCBI accession numbers provided in the figure) was performed using the EBI web based Clustal Omega tool (doi: 10.1093/nar/gkz268). The phylogenetic tree was built with the “one click” mode for phylogenetic analysis (http://www.phylogeny.fr/) with standard settings (doi: 10.1093/nar/gkn180; doi: 10.1186/1471-2148-10-8).

#### RNA isolation from primary tissues

Sections of liver and spleen were preserved in RNAlater Stabilization Solution (Thermo Fisher Scientific) and frozen. Thawed samples were transferred to PBS and homogenized. One-eighth of the homogenate was mixed with TRIzol Reagent (Thermo Fisher Scientific). BMDM were directly lysed in TRIzol Reagent and frozen. Total RNA was isolated according to the manufacturer’s protocol. Liver and spleen RNA were subject to DNase treatment (TURBO DNA-Free kit, Thermo Fisher Scientific) per the manufacturer’s protocol prior to RNA-Seq analysis. BMDM RNA was analyzed by one-step qRT-PCR using QuantiFast SYBR Green RT-PCR Kit (Qiagen) using Applied Biosciences 7500 Fast Real-Time PCR System.

#### RNA library construction and sequencing

The quantity and quality of the extracted RNA was assessed using a Thermo Fisher Scientific qubit 2.0 fluorometer with the Qubit RNA BR Assay Kit (Thermo Fisher Scientific, Q10211) and an Advanced Analytical Fragment Analyzer System using a Fragment Analyzer RNA Kit (Agilent, DNF-471), respectively. Sequencing libraries were prepared using an Illumina TruSeq Stranded Total RNA Library Prep Gold kit (Illumina, 20020599) in combination with TruSeq RNA UD Indexes (Illumina, 20022371) according to the Illumina guidelines. Sequencing libraries were sequenced paired-end (2 × 50 bp) using an Illumina NovaSeq 6000 S1 Reagent Kit (100 cycles; Illumina, 20012865) on an Illumina NovaSeq 6000 sequencer. The quality control assessments, generation of libraries and sequencing run were all performed at the Next Generation Sequencing Platform, University of Bern, Switzerland.

Data generated from individual samples (>30 million read per sample, single read 50-mers) were mapped separately against the GRCm38 murine reference genome. Gene expression was calculated for individual transcripts as reads per kilobase per million bases mapped (RPKM). All transcriptomic analyses were performed using CLC Genomics Workbench 20 (Qiagen, Aarhaus).

Differentially expressed genes (DEGs) were identified by calculating fold changes in expression, p-values were corrected by taking false discovery rate (FDR) for multiple comparison into account. Mus Musculus EBI Gene Ontology Annotation Database was used to execute Gene ontology (GO) Enrichment Analyses for biological processes. Gene identifiers for DEGs absolute FC > 5, RPKM > 2) were used as input and identification of significantly enriched GO categories. P-values for specific GO categories were generated after Bonferroni correction for multiple testing. Z-scores, used as indicator for activation of biological processes (positive value) or inactivation (negative value), were calculated based on the expression fold change as follows:

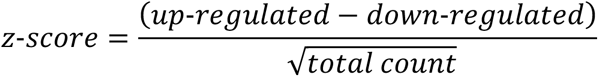

#### Antibodies for immunofluorescence and flow cytometry

HCoV-229E infection was detected by staining permeabilized cells for N protein (Anticuerpo Monoclonal, 1E7, Ingenasa) and a secondary donkey anti-mouse AlexaFluor488-conjugated antibody (Jackson Immuno Research). HCoV-OC43 infection was detected by staining permeabilized cells for nucleoprotein (MAB9013, Millipore) and a secondary goat anti-mouse AlexaFluor488-conjugated antibody (Thermo Fisher Scientific). ZIKV infected cells were stained for viral antigen using a monoclonal antibody anti-flavivirus group antigen 4G2 (MAB10216; RRID: AB_827205, EMD Millipore) as primary antibody and AlexaFluor488 (A-11001, RRID: AB_2534069, Thermo Fisher Scientific) as secondary antibody. CD13 was stained anti-human CD13 Antibody, APC conjugated (clone WM15, BioLegend). The following antibodies were used to differentiate immune cell populations: anti-CD3ε-PE (145-2C11, Tonbo Biosciences), anti-CD3ε-FITC (145-2C11, Tonbo Biosciences), anti-CD11c-PECy7 (N418, Tonbo Biosciences), anti-Ly6G-PE (1A8, Tonbo Biosciences), anti-Ly6G-FITC (1A8, Tonbo Biosciences), anti-Ly6g-PerCPCy5.5 (1A8, Tonbo Biosciences), anti-CD11b-PECy7 (M1/70, Tonbo Biosciences), anti-CD19-PE (6D5, BioLegend), anti-CD19-FITC (1D3, Tonbo Biosciences), anti-CD19-PECy5 (6D5, BioLegend), anti-F4/80-PECy5 (BM8, BioLegend), anti-Nkp46-PECy7 (29A1.4, BioLegend), anti-CD4-PECy5 (GK1.5, Tonbo Biosciences), anti-CD8-PECy7 (53-6.7, Tonbo Biosciences).

#### Primers and gBlocks

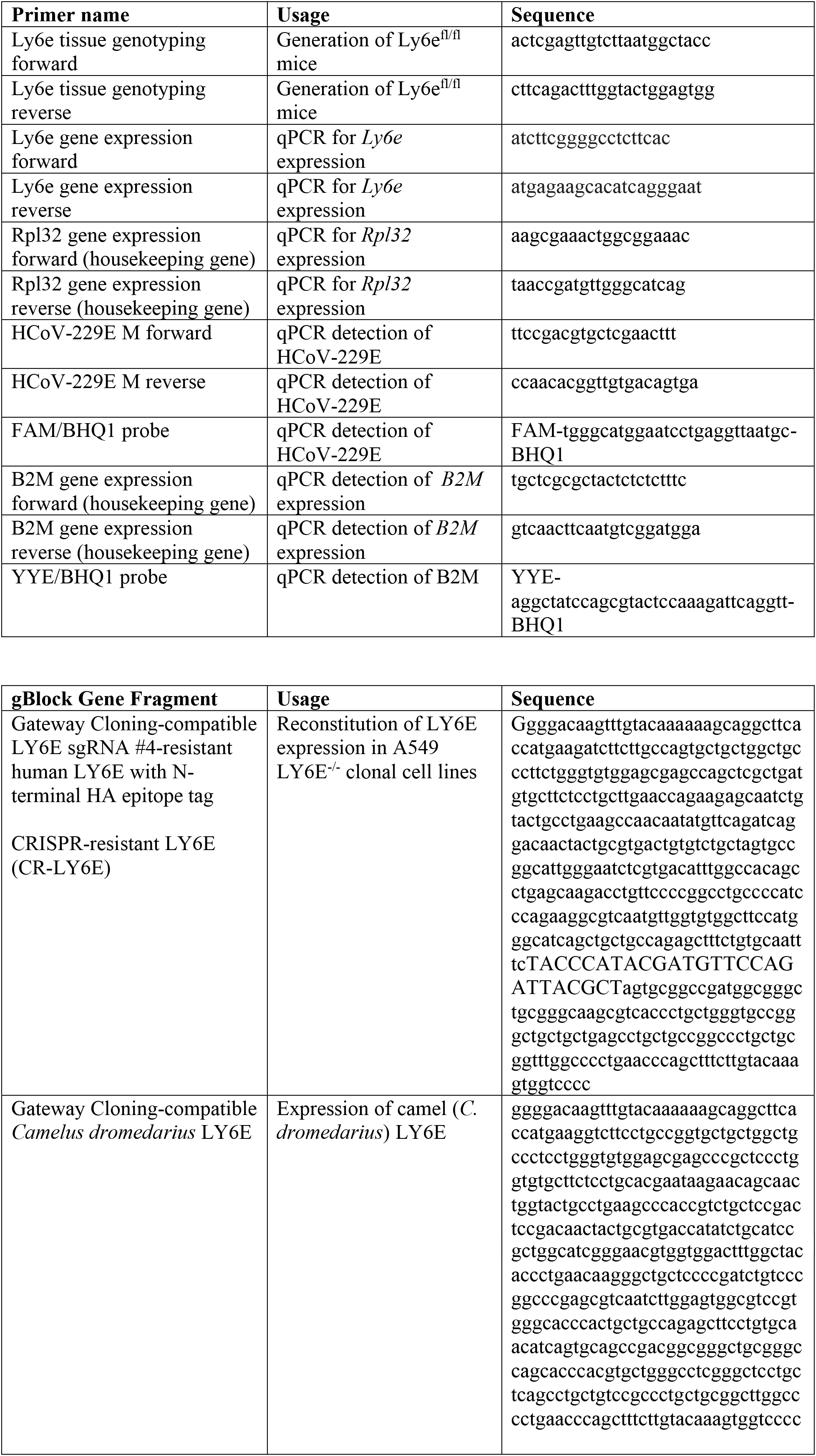

## References

1 Zhu, N. et al. A Novel Coronavirus from Patients with Pneumonia in China, 2019. N Engl J Med, doi:10.1056/NEJMoa2001017 (2020).

2 Kindler, E., Thiel, V. & Weber, F. Interaction of SARS and MERS Coronaviruses with the Antiviral Interferon Response. Adv Virus Res 96, 219–243, doi:10.1016/bs.aivir.2016.08.006 (2016).

3 Corman, V. M., Muth, D., Niemeyer, D. & Drosten, C. Hosts and Sources of Endemic Human Coronaviruses. Adv Virus Res 100, 163–188, doi:10.1016/bs.aivir.2018.01.001 (2018).

4 Wang, B. X. & Fish, E. N. Global virus outbreaks: Interferons as 1st responders. Semin Immunol 43, 101300, doi:10.1016/j.smim.2019.101300 (2019).

5 Cervantes-Barragan, L. et al. Control of coronavirus infection through plasmacytoid dendritic- cell-derived type I interferon. Blood 109, 1131–1137, doi:10.1182/blood-2006-05-023770 (2007).

6 Schoggins, J. W. Interferon-Stimulated Genes: What Do They All Do? Annu Rev Virol 6, 567–584, doi:10.1146/annurev-virology-092818-015756 (2019).

7 Schoggins, J. W. et al. A diverse range of gene products are effectors of the type I interferon antiviral response. Nature 472, 481–485, doi:10.1038/nature09907 (2011).

8 Dittmann, M. et al. A serpin shapes the extracellular environment to prevent influenza A virus maturation. Cell 160, 631–643, doi:10.1016/j.cell.2015.01.040 (2015).

9 Mar, K. B. et al. LY6E mediates an evolutionarily conserved enhancement of virus infection by targeting a late entry step. Nature Communications 9, 3603, doi:10.1038/s41467-018-06000-y (2018).

10 Hackett, B. A. & Cherry, S. Flavivirus internalization is regulated by a size-dependent endocytic pathway. Proc Natl Acad Sci U S A 115, 4246–4251, doi:10.1073/pnas.1720032115 (2018).

11 Yu, J., Liang, C. & Liu, S. L. Interferon-inducible LY6E Protein Promotes HIV-1 Infection. J Biol Chem 292, 4674–4685, doi:10.1074/jbc.M116.755819 (2017).

12 Yu, J., Liang, C. & Liu, S. L. CD4-Dependent Modulation of HIV-1 Entry by LY6E. J Virol 93, doi:10.1128/JVI.01866-18 (2019).

13 Sikkema, R. S. et al. Global status of Middle East respiratory syndrome coronavirus in dromedary camels: a systematic review. Epidemiol Infect 147, e84, doi:10.1017/S095026881800345X (2019).

14 Upadhyay, G. Emerging Role of Lymphocyte Antigen-6 Family of Genes in Cancer and Immune Cells. Front Immunol 10, 819, doi:10.3389/fimmu.2019.00819 (2019).

15 Heald-Sargent, T. & Gallagher, T. Ready, set, fuse! The coronavirus spike protein and acquisition of fusion competence. Viruses 4, 557–580, doi:10.3390/v4040557 (2012).

16 Park, J. E. et al. Proteolytic processing of Middle East respiratory syndrome coronavirus spikes expands virus tropism. Proc Natl Acad Sci U S A 113, 12262–12267, doi:10.1073/pnas.1608147113 (2016).

17 Kleine-Weber, H., Elzayat, M. T., Hoffmann, M. & Pohlmann, S. Functional analysis of potential cleavage sites in the MERS-coronavirus spike protein. Sci Rep 8, 16597, doi:10.1038/s41598-018-34859-w (2018).

18 Langford, M. B., Outhwaite, J. E., Hughes, M., Natale, D. R. C. & Simmons, D. G. Deletion of the Syncytin A receptor Ly6e impairs syncytiotrophoblast fusion and placental morphogenesis causing embryonic lethality in mice. Sci Rep 8, 3961, doi:10.1038/s41598-018-22040-2 (2018).

19 Matthews, A. E. et al. Antibody is required for clearance of infectious murine hepatitis virus A59 from the central nervous system, but not the liver. J Immunol 167, 5254–5263, doi:10.4049/jimmunol.167.9.5254 (2001).

20 Phares, T. W. et al. CD4 T cells promote CD8 T cell immunity at the priming and effector site during viral encephalitis. J Virol 86, 2416–2427, doi:10.1128/JVI.06797-11 (2012).

21 Wijburg, O. L., Heemskerk, M. H., Sanders, A., Boog, C. J. & Van Rooijen, N. Role of virus-specific CD4+ cytotoxic T cells in recovery from mouse hepatitis virus infection. Immunology 87, 34–41 (1996).

22 Wijburg, O. L., Heemskerk, M. H., Boog, C. J. & Van Rooijen, N. Role of spleen macrophages in innate and acquired immune responses against mouse hepatitis virus strain A59. Immunology 92, 252–258, doi:10.1046/j.1365-2567.1997.00340.x (1997).

23 Cervantes-Barragan, L. et al. Type I IFN-mediated protection of macrophages and dendritic cells secures control of murine coronavirus infection. J Immunol 182, 1099–1106 (2009).

24 Weiss, S. R. & Leibowitz, J. L. Coronavirus pathogenesis. Adv Virus Res 81, 85–164, doi:10.1016/B978-0-12-385885-6.00009-2 (2011).

25 Eriksson, K. K., Cervantes-Barragan, L., Ludewig, B. & Thiel, V. Mouse hepatitis virus liver pathology is dependent on ADP-ribose-1’’-phosphatase, a viral function conserved in the alpha-like supergroup. J Virol 82, 12325–12334, doi:10.1128/JVI.02082-08 (2008).

26 Karnam, G. et al. CD200 receptor controls sex-specific TLR7 responses to viral infection. PLoS Pathog 8, e1002710, doi:10.1371/journal.ppat.1002710 (2012).

27 Hong, S. et al. B Cells Are the Dominant Antigen-Presenting Cells that Activate Naive CD4(+) T Cells upon Immunization with a Virus-Derived Nanoparticle Antigen. Immunity 49, 695–708 e694, doi:10.1016/j.immuni.2018.08.012 (2018).

28 Mempel, T. R., Henrickson, S. E. & Von Andrian, U. H. T-cell priming by dendritic cells in lymph nodes occurs in three distinct phases. Nature 427, 154–159, doi:10.1038/nature02238 (2004).

29 Yu, J. & Liu, S. L. Emerging Role of LY6E in Virus-Host Interactions. Viruses 11, doi:10.3390/v11111020 (2019).

30 Lalezari, J. P. et al. Enfuvirtide, an HIV-1 fusion inhibitor, for drug-resistant HIV infection in North and South America. N Engl J Med 348, 2175–2185, doi:10.1056/NEJMoa035026 (2003)

## References

31 Thiel, V. & Siddell, S. G. Reverse genetics of coronaviruses using vaccinia virus vectors. Curr Top Microbiol Immunol 287, 199–227, doi:10.1007/3-540-26765-4_7 (2005).

32 van den Worm, S. H. et al. Reverse genetics of SARS-related coronavirus using vaccinia virus-based recombination. PLoS One 7, e32857, doi:10.1371/journal.pone.0032857 (2012).

33 Bruckova, M., McIntosh, K., Kapikian, A. Z. & Chanock, R. M. The adaptation of two human coronavirus strains (OC38 and OC43) to growth in cell monolayers. Proc Soc Exp Biol Med 135, 431–435, doi:10.3181/00379727-135-35068 (1970).

34 Coley, S. E. et al. Recombinant mouse hepatitis virus strain A59 from cloned, full-length cDNA replicates to high titers in vitro and is fully pathogenic in vivo. J Virol 79, 3097–3106, doi:10.1128/JVI.79.5.3097-3106.2005 (2005).

35 Lundin, A. et al. Targeting membrane-bound viral RNA synthesis reveals potent inhibition of diverse coronaviruses including the middle East respiratory syndrome virus. PLoS Pathog 10, e1004166, doi:10.1371/journal.ppat.1004166 (2014).

36 Kindler, E. et al. Efficient replication of the novel human betacoronavirus EMC on primary human epithelium highlights its zoonotic potential. mBio 4, e00611–00612, doi:10.1128/mBio.00611-12 (2013).

37 Thiel, V. et al. Mechanisms and enzymes involved in SARS coronavirus genome expression. J Gen Virol 84, 2305–2315, doi:10.1099/vir.0.19424-0 (2003).

38 Richardson, R. B. et al. A CRISPR screen identifies IFI6 as an ER-resident interferon effector that blocks flavivirus replication. Nat Microbiol 3, 1214–1223, doi:10.1038/s41564-018-0244-1 (2018).

39 Berger Rentsch, M. & Zimmer, G. A vesicular stomatitis virus replicon-based bioassay for the rapid and sensitive determination of multi-species type I interferon. PLoS One 6, e25858, doi:10.1371/journal.pone.0025858 (2011).

40 Zhang, L. et al. Infection of ciliated cells by human parainfluenza virus type 3 in an in vitro model of human airway epithelium. J Virol 79, 1113–1124, doi:10.1128/JVI.79.2.1113-1124.2005 (2005).

41 Biacchesi, S. et al. Recovery of human metapneumovirus from cDNA: optimization of growth in vitro and expression of additional genes. Virology 321, 247–259, doi:10.1016/j.virol.2003.12.020 (2004).

42 Jones, C. T. et al. Real-time imaging of hepatitis C virus infection using a fluorescent cell-based reporter system. Nat Biotechnol 28, 167–171, doi:10.1038/nbt.1604 (2010).

43 Schoggins, J. W. et al. Dengue reporter viruses reveal viral dynamics in interferon receptor-deficient mice and sensitivity to interferon effectors in vitro. Proc Natl Acad Sci U S A 109, 14610–14615, doi:10.1073/pnas.1212379109 (2012).

44 McGee, C. E. et al. Infection, dissemination, and transmission of a West Nile virus green fluorescent protein infectious clone by Culex pipiens quinquefasciatus mosquitoes. Vector Borne Zoonotic Dis 10, 267–274, doi:10.1089/vbz.2009.0067 (2010).

45 Liu, D. et al. Fast hepatitis C virus RNA elimination and NS5A redistribution by NS5A inhibitors studied by a multiplex assay approach. Antimicrob Agents Chemother 59, 3482–3492, doi:10.1128/AAC.00223-15 (2015).

46 Suthar, M. S., Shabman, R., Madric, K., Lambeth, C. & Heise, M. T. Identification of adult mouse neurovirulence determinants of the Sindbis virus strain AR86. J Virol 79, 4219–4228, doi:10.1128/JVI.79.7.4219-4228.2005 (2005).

47 Petrakova, O. et al. Noncytopathic replication of Venezuelan equine encephalitis virus and eastern equine encephalitis virus replicons in Mammalian cells. J Virol 79, 7597–7608, doi:10.1128/JVI.79.12.7597-7608.2005 (2005).

48 Tsetsarkin, K. et al. Infectious clones of Chikungunya virus (La Reunion isolate) for vector competence studies. Vector Borne Zoonotic Dis 6, 325–337, doi:10.1089/vbz.2006.6.325 (2006).

49 Hoffmann, H. H. et al. Diverse Viruses Require the Calcium Transporter SPCA1 for Maturation and Spread. Cell Host Microbe 22, 460–470 e465, doi:10.1016/j.chom.2017.09.002 (2017).

50 Torriani, G. et al. Identification of Clotrimazole Derivatives as Specific Inhibitors of Arenavirus Fusion. J Virol 93, doi:10.1128/JVI.01744-18 (2019).

51 Gorisek, J. & Marzan, B. [Changes in the blood picture and blood coagulation in calf poisoned by the fern Pteridium aquilinum]. Wien Tierarztl Monatsschr 52, 530–538 (1965).

52 Gierer, S. et al. The spike protein of the emerging betacoronavirus EMC uses a novel coronavirus receptor for entry, can be activated by TMPRSS2, and is targeted by neutralizing antibodies. J Virol 87, 5502–5511, doi:10.1128/JVI.00128-13 (2013).

53 Hoffmann, M. et al. Differential sensitivity of bat cells to infection by enveloped RNA viruses: coronaviruses, paramyxoviruses, filoviruses, and influenza viruses. PLoS One 8, e72942, doi:10.1371/journal.pone.0072942 (2013).

54 Qing, E., Hantak, M., Perlman, S. & Gallagher, T. Distinct Roles for Sialoside and Protein Receptors in Coronavirus Infection. mBio 11, doi:10.1128/mBio.02764-19 (2020).

55 Hoffmann, M. et al. Fusion-active glycoprotein G mediates the cytotoxicity of vesicular stomatitis virus M mutants lacking host shut-off activity. J Gen Virol 91, 2782–2793, doi:10.1099/vir.0.023978-0 (2010).

56 Locher, S., Schweneker, M., Hausmann, J. & Zimmer, G. Immunogenicity of propagation-restricted vesicular stomatitis virus encoding Ebola virus glycoprotein in guinea pigs. J Gen Virol 99, 866–879, doi:10.1099/jgv.0.001085 (2018).

57 Hanika, A. et al. Use of influenza C virus glycoprotein HEF for generation of vesicular stomatitis virus pseudotypes. J Gen Virol 86, 1455–1465, doi:10.1099/vir.0.80788-0 (2005).

58 Skarnes, W. C. et al. A conditional knockout resource for the genome-wide study of mouse gene function. Nature 474, 337–342, doi:10.1038/nature10163 (2011).

59 Schickli, J. H., Thackray, L. B., Sawicki, S. G. & Holmes, K. V. The N-terminal region of the murine coronavirus spike glycoprotein is associated with the extended host range of viruses from persistently infected murine cells. J Virol 78, 9073–9083, doi:10.1128/JVI.78.17.9073-9083.2004 (2004).

60 Agac, D., Estrada, L. D., Maples, R., Hooper, L. V. & Farrar, J. D. The beta2-adrenergic receptor controls inflammation by driving rapid IL-10 secretion. Brain Behav Immun 74, 176–185, doi:10.1016/j.bbi.2018.09.004 (2018).

61 Michailidis, E. et al. A robust cell culture system supporting the complete life cycle of hepatitis B virus. Sci Rep 7, 16616, doi:10.1038/s41598-017-16882-5 (2017).

62 Widjaja, I. et al. Towards a solution to MERS: protective human monoclonal antibodies targeting different domains and functions of the MERS-coronavirus spike glycoprotein. Emerg Microbes Infect 8, 516–530, doi:10.1080/22221751.2019.1597644 (2019).

63 Vijgen, L. et al. Development of one-step, real-time, quantitative reverse transcriptase PCR assays for absolute quantitation of human coronaviruses OC43 and 229E. J Clin Microbiol 43, 5452–5456, doi:10.1128/JCM.43.11.5452-5456.2005 (2005).

